# TBK1-mediated phosphorylation of LC3C and GABARAP-L2 controls autophagosome shedding by ATG4 protease

**DOI:** 10.1101/634519

**Authors:** Lina Herhaus, Ramachandra M. Bhaskara, Alf Håkon Lystad, Anne Simonsen, Gerhard Hummer, Ivan Dikic

## Abstract

Autophagy is a highly conserved catabolic process through which defective or otherwise harmful cellular components are targeted for degradation via the lysosomal route. Regulatory pathways, involving post-translational modifications such as phosphorylation, play a critical role in controlling this tightly orchestrated process. Here, we demonstrate that TBK1 regulates autophagy by phosphorylating autophagy modifiers LC3C and GABARAP-L2 on surface-exposed serine residues (LC3C S93 and S96; GABARAP-L2 S87 and S88). This phosphorylation event impedes their binding to the processing enzyme ATG4 by destabilizing the complex. Phosphorylated LC3C/GABARAP-L2 cannot be removed from liposomes by ATG4 and are thus protected from ATG4-mediated premature removal from nascent autophagosomes. This ensures a steady coat of lipidated LC3C/GABARAP-L2 throughout the early steps in autophagosome formation and aids in maintaining a unidirectional flow of the autophagosome to the lysosome. Taken together, we present a new regulatory mechanism of autophagy, which influences the conjugation and de-conjugation of LC3C and GABARAP-L2 to autophagosomes by TBK1-mediated phosphorylation.

## Introduction

The recycling of redundant cytosolic components and damaged organelles is termed autophagy. It is a highly conserved process, which increases during starvation conditions or during other cellular stresses (Dikic, 2017, Xie & Zhou, 2018). Autophagy involves the formation of the phagophore, a double-membrane cup-shaped structure, which expands to enwrap and enclose the designated cellular cargo to form the autophagosome, which then fuses with lysosomes to enable enzymatic degradation of its cargo along with its inner membrane (Yang & Klionsky, 2010). Upon induction of autophagy, small ubiquitin-like LC3 proteins (autophagy-modifiers) are conjugated to phosphatidylethanolamine (PE) anchoring them to the growing phagophore membrane. This conjugation is carried out by the lipidation cascade enzymes (ATG3, ATG5, ATG7, ATG12, ATG16L1) and allows cargo selection and autophagosome formation (Nakatogawa, 2013, Stolz et al., 2014). In humans, there are six autophagy-modifier proteins, grouped into two sub-families: (1) LC3A, LC3B, LC3C, and (2) GABARAP, GABARAP-L1, and GABARAP-L2/GATE-16 (Cemma et al., 2016).

LC3 proteins undergo two processing steps, (1) an initial proteolytic cleavage of the peptide bond responsible for the conversion of pro-LC3 to active LC3 and (2) the subsequent cleavage of the amide bond for de-lipidation of LC3-PE from autophagosomes to regenerate a free cytosolic LC3-pool. Both processing steps are catalyzed by ATG4 (Zhang et al., 2016). Among the four mammalian paralogs of ATG4, ATG4B is the most active protease followed by ATG4A and ATG4C/D, which exhibit minimal protease activity (Li et al., 2011). LC3s are bound by the ATG4B enzyme body and through LC3 interacting motifs (LIRs) located at the N- and C-terminal flexible tails of ATG4B (Maruyama & Noda, 2017). ATG4s are cysteine proteases that cleave peptide bonds of pro-LC3 to expose the C-terminal glycine and allow conjugation with PE. ATG4 also de-conjugates LC3-PE from the outer membrane of autophagosomes preceding, or just after autophagosome-lysosome fusion by cleaving the amide bond between PE and the C-terminal glycine residue of LC3s (Yu et al., 2012). Both processing steps are known to be regulated by direct phosphorylation of ATG4B itself. Phosphorylation of ATG4B at S383/392 increases its protease activity (Yang et al., 2015), especially during the LC3 delipidation phase, whereas ULK1-mediated phosphorylation of ATG4B at S316 (in humans) (Pengo et al., 2017) or at S307 (in yeast) (Sanchez-Wandelmer et al., 2017) reduces pro-LC3 binding and C-terminal tail cleavage. Likewise, oxidation of ATG4 by H_2_O_2_ also attenuates its activity and blocks LC3 de-lipidation (Scherz-Shouval et al., 2007).

The serine-threonine kinase TBK1 has been implicated in the selective degradation of depolarized mitochondria (mitophagy) and intracellular pathogens (xenophagy) (Randow & Youle, 2014, Richter et al., 2016b, Wild et al., 2011). Specific autophagic cargo marked with an ubiquitin signal is recognized by autophagy receptor proteins such as Optineurin (OPTN) and p62 (Herhaus & Dikic, 2015, Herhaus & Dikic, 2018). They physically bridge the cargo to the nascent phagophore by binding to ubiquitin via their UBD and to LC3 family proteins via their conserved LIR motifs, respectively. These autophagy receptors also recruit TBK1 to the site of autophagosome formation. Local accumulation of active TBK1 phosphorylates p62 and OPTN, which increases their binding affinity for polyubiquitin chains and the LC3 family proteins, thereby driving autophagy (Heo et al., 2015, Lazarou et al., 2015, Matsumoto et al., 2011, Ordureau et al., 2015, Pilli et al., 2012, Richter et al., 2016a, Thurston et al., 2009, Wild et al., 2011).

In this study we expand the role of TBK1 in the autophagic process by demonstrating its ability to directly phosphorylate LC3C on S93/96 and GABARAP-L2 on S87/88. We study the consequences of LC3 phosphorylation during autophagy and show that this phosphorylation primarily impedes ATG4-mediated processing of LC3 on the liposomes, adding a new layer of regulation.

## Results

### TBK1 phosphorylates LC3C and GABARAP-L2 *in vitro*

The serine-threonine kinase TBK1 has previously been shown to phosphorylate autophagy receptors such as OPTN and p62 (Pilli et al., 2012, Richter et al., 2016a, Wild et al., 2011). To test if recombinant TBK1 can also directly phosphorylate autophagy modifiers, we performed an *in vitro* kinase assay. Four out of the six autophagy modifier proteins: LC3A, LC3C, GABARAP-L1 and GABARAP-L2 are directly phosphorylated by TBK1 *in vitro* (Figure 1A). The S/T phosphorylation sites of LC3-family proteins were identified by mass spectrometry with significant PEP scores (Figure 1B) and we decided to further investigate the TBK1-mediated phosphorylation sites of LC3C at S93 and S96 and GABARAP-L2 at S87 and S88 in detail. The TBK1-mediated phosphorylation sites of LC3C (Figure 1C) and GABARAP-L2 (Figure 1D) are topologically equivalent and are present in surface exposed loops (depicted in red). This loop is on the opposite face of the LIR binding pocket indicating that LIR-mediated interactions of LC3C might not be affected directly upon phosphorylation.

**Figure 1:**
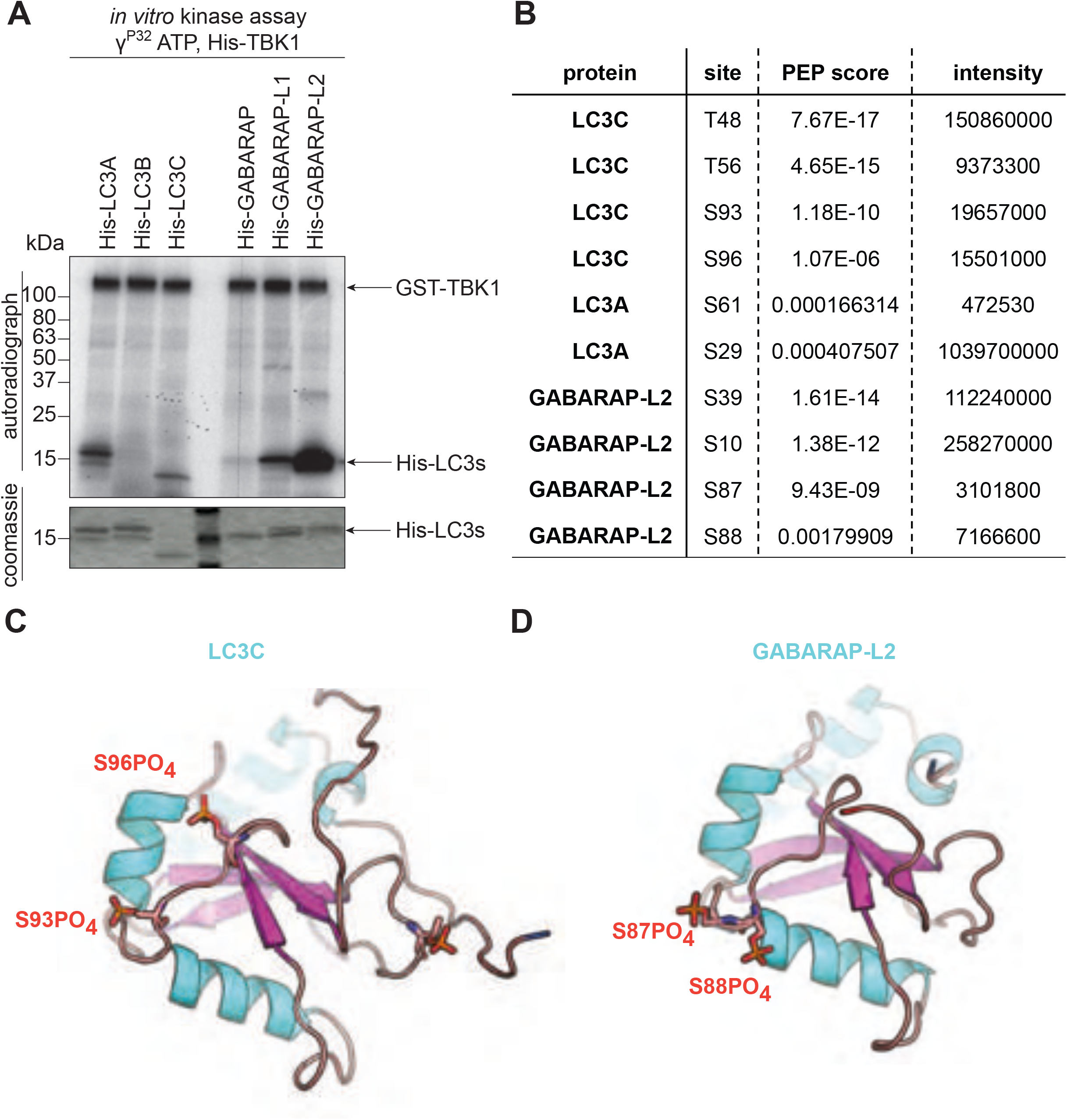
TBK1 phosphorylates LC3C and GABARAP-L2 *in vitro*. (A) Coomassie stain and autoradiography of SDS-PAGE after an *in vitro* kinase assay with His-TBK1 and His-LC3 family proteins as substrates. TBK1 phosphorylates LC3A, LC3C, GABARAP-L1 and GABARAP-L2 *in vitro.* (B) Identification of phosphosites by mass spectrometry following an *in vitro* TBK1 kinase assay. (C) Structure of LC3C_8-125_ (PDB: 3WAM) with modeled phosphate groups (red sticks) at S93 and S96. (D) Topologically equivalent positions in GABARAP-L2 (PDB: 4CO7), S87 and S88 are also phosphorylated (red sticks) by TBK1. Phosphorylation sites are on the opposite face of the LIR binding pocket of LC3-proteins.

### TBK1 phosphorylates and binds LC3C and GABARAP-L2 *in cells*

To test if TBK1 also phosphorylates LC3C in cells, HEK293T cells were SILAC labeled and either WT TBK1 (heavy label) or TBK1 kinase dead (K38A; light label) were overexpressed along with GFP-LC3C or with GFP-GABARAP-L2. GFP-proteins were immunoprecipitated and analyzed by mass spectrometry. Phosphorylation at positions S96 and S93 of LC3C was enhanced in the presence of WT TBK1 (factors 6 and 10 respectively; Figure 2A), as compared to TBK1 K38A. Similarly, the presence of WT TBK1 resulted in enhanced phosphorylation of GABARAP-L2 at S87 and S88 (by factors 13 and 2, respectively; Figure 2A). Unfortunately, our efforts to generate phospho-specific antibodies against GABARAP-L2 S87-PO_4_ and GABARAP-L2 S87/88-PO_4_ failed (Figure S1). To confirm the phosphorylation event directly, we visualized it by using phos-tag^TM^ polyacrylamide gels, where phosphorylated proteins are retained by the phos-tag reagent and appear at a higher molecular weight. Overexpression of WT TBK1, but not TBK1 kinase dead, induced an upward shift and retention of phosphorylated LC3C (Figure 2B). The ratio of phosphorylated to unphosphorylated LC3C is also higher upon the induction of mitophagy, by the addition of Parkin, an E3 ligase and CCCP, the mitochondrial depolarization agent (compare lanes two and five of phosphorylated upper HA-band from phos-tag gel, Figure 2B). Moreover, the endogenous TBK1 from HeLa or HEK293T cell lysate binds to GST-LC3C and GST-GABARAP-L2 (Figure 2C) indicating direct physical interaction. The binding of TBK1 to LC3C and GABARAP-L2 is independent of its catalytic activity (Figure 2D) and could be mediated through its C-terminal coiled-coil region (Figure 2D), which is known to bind OPTN (Freischmidt et al., 2015).

**Figure 2:**
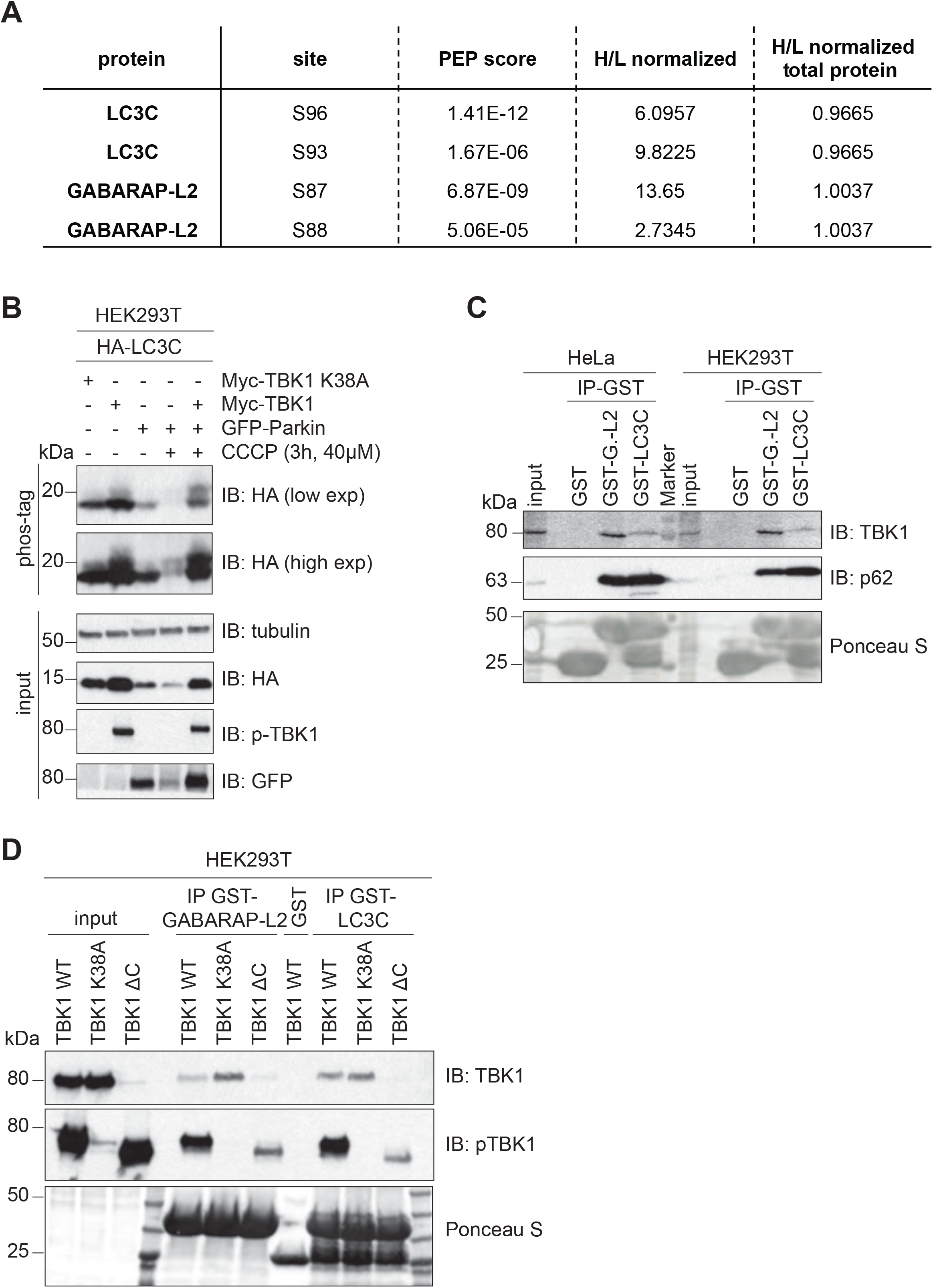
TBK1 phosphorylates and binds LC3C and GABARAP-L2 *in cells*. (A) Identification of phosphosites by mass spectrometry following GFP-LC3C or GFP-GABARAP-L2 immunoprecipitation (IP) in SILAC labeled cells. TBK1 WT was overexpressed in heavy and TBK1 kinase dead K38A was overexpressed in light labeled cells. (B) SDS-PAGE and Western blot of phos-tag^TM^ gel with HEK293T cell lysates. Cells were transfected with HA-LC3C, TBK1 WT or K38A and GFP-Parkin and left untreated or treated with CCCP (3 hours, 40 μM) to induce mitophagy. (C) SDS-PAGE and Western blot of HEK293T and HeLa cell lysates and GST-LC3C or GST-GABARAP-L2 immunoprecipitations. (D) SDS-PAGE and Western blot of HEK293T cell lysates transfected with full-length TBK1, a C-terminal truncation mutant (TBK1 ΔC), a kinase dead version (TBK1 K38A) and GST-LC3C or GST-GABARAP-L2 immunoprecipitations.

### Phospho-mimetic LC3C impedes ATG4 cleavage and binding

To understand the consequences of phosphorylated-LC3C, we looked at its phospho-sites in more detail. The LC3C phosphorylation sites S93 and S96 are situated on the face opposite to the hydrophobic pocket enabling LIR binding (Figure 1C) and are therefore less likely to influence the direct binding of LC3C to autophagy receptors or adaptors. However, they are in close proximity to the C-terminal tail of LC3C which is proteolytically processed. ATG4 mediated processing of the LC3C C-terminal tail allows lipid-conjugation and adherence to autophagosomes. To test if phosphorylation of these residues could impair the proteolytic cleavage of the LC3C C-terminal tail by ATG4, an *in vitro* cleavage assay was performed. Double-tagged LC3C WT, S93/96A or phospho-mimetic LC3C S93/96D were incubated with ATG4B for indicated times and the C-terminal cleavage of LC3C was monitored by detecting the appearance of truncated His-LC3C protein (Figure 3A). ATG4B cleaves the entire pool of LC3C WT or S93/96A within 10 minutes, whereas only half of the phospho-mimetic LC3C S93/96D pool is cleaved (Figure 3A). When LC3 proteins are overexpressed in HEK293T cells, they are rapidly processed by endogenous ATG4 proteins. The C-terminal tail of LC3C that is cleaved by ATG4s is considerably larger (21 residues) than that of other LC3 family proteins. Hence, a pro-form of LC3C S93/96D could be visualized by separating cell lysate on a 15% polyacrylamide gel (Figure 3B).

**Figure 3:**
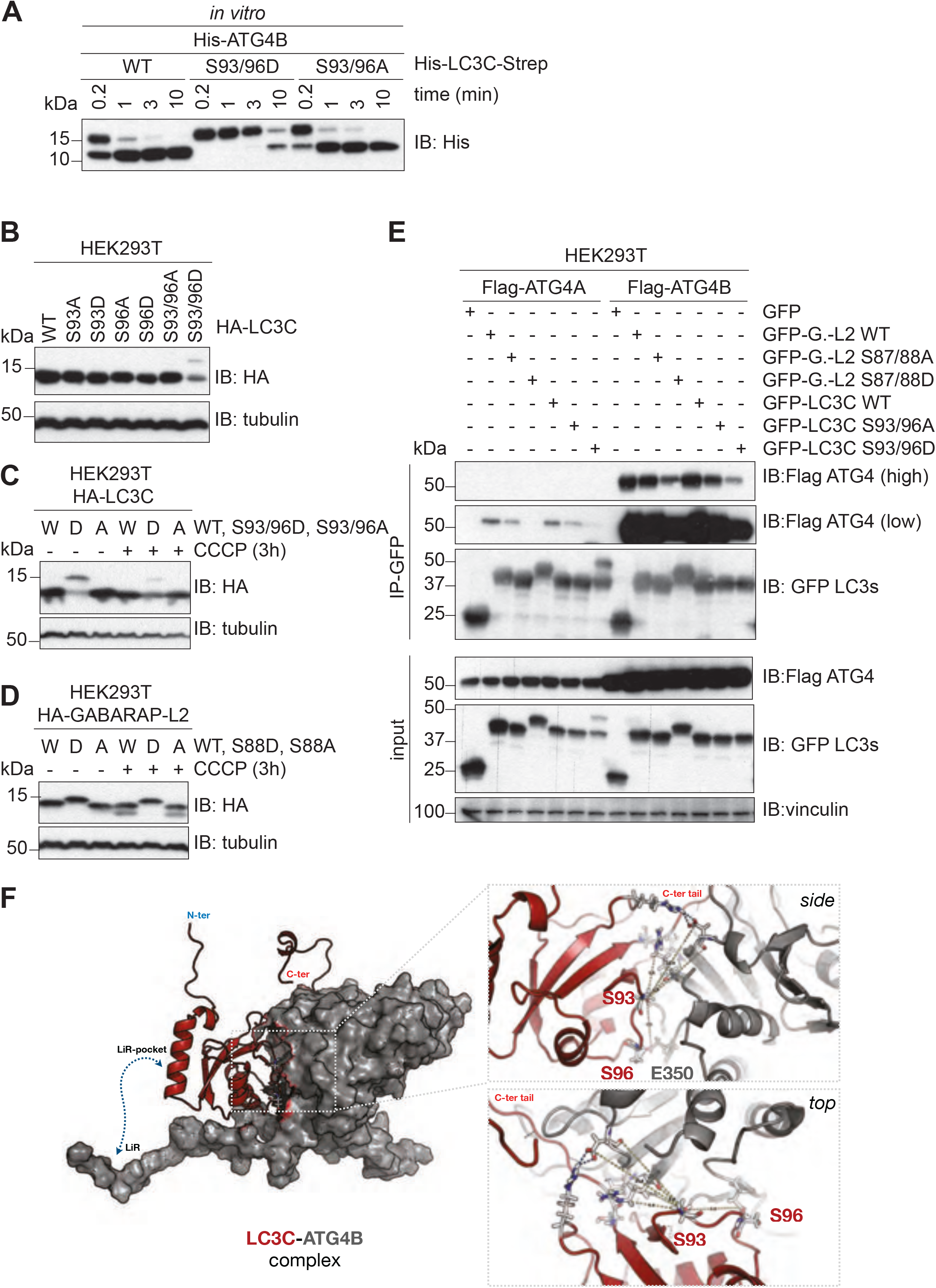
Phospho-mimetic LC3C and GABARAP-L2 impede ATG4 cleavage and binding. (A) SDS-PAGE and Western blotting of *in vitro* ATG4 cleavage assay. Purified double tagged His-LC3C-Strep WT and mutants were incubated with ATG4B for indicated time points. LC3C S93/96D mutation slows down C-terminal cleavage of LC3C by ATG4B. (B) SDS-PAGE and Western blot of HEK293T cell lysates transfected with LC3C WT or mutants. S93/96D mutation of LC3C impedes cleavage of pro-LC3 by endogenous ATG4s. (C,D) SDS-PAGE and Western blot of HEK293T cell lysates transfected with LC3C WT or mutants (C) or GABARAP-L2 WT or mutants (D). Cells were left untreated or treated with 40 µM CCCP for 3 hours to induce mitophagy. (E) SDS-PAGE and Western blot of HEK293T cell lysates and GFP immunoprecipitations. Cells were transfected with Flag-tagged ATG4A or ATG4B and GFP-tagged LC3C or GABARAP-L2 WT or mutants and lysates used for GFP IPs. S93/96D mutation of LC3C and S87/88D mutation of GABARAP-L2 impede binding to ATG4A or B. (F) Full-length LC3C (red cartoon) binds to ATG4B (grey surface) with its C-terminal tail accessible to the active site of ATG4B. Phosphorylation of LC3C at S93 and S96 (sticks) affect binding to ATG4B. Zoom-up showing the side and top view of LC3C-ATG4B interface. S96 of LC3C and E350 of ATG4B form direct hydrogen-bonds across the interface. S93 position is central to a network of polar interactions (blue dashed lines; side chains shown as sticks) across the interface.

The inability of ATG4 to process phosphorylated LC3C might be distinct during stress conditions. To test this, we induced mitophagy in HEK293T cells by adding CCCP. Upon induction of mitophagy LC3C S93/96D could not be completely processed by ATG4s (Figure 3C). Similarly, GABARAP-L2 phospho-mimetic (S88D) could not be cleaved by endogenous ATG4s, impairing subsequent lipidation (Figure 3D). We reasoned that this inability of ATG4 to process LC3C S93/96D and GABARAP-L2 S87/88D could be due to an impediment in direct protein binding, and therefore tested this by co-expressing GFP-LC3 proteins with either Flag-ATG4A or Flag-ATG4B in HEK293T cells and subjected them to GFP-immunoprecipitation (Figure 3E). Phospho-mimetic mutants LC3C S93/96D and GABARAP-L2 S87/88D displayed reduced binding to ATG4A and B. To understand this reduced binding, we modelled the full-length LC3C-ATG4B complex based on the core crystal structure (Satoo et al., 2009) (see Methods). We tested the effect of phosphorylation at both these sites (S93 and S96) by modeling phosphate groups onto serine residues in the LC3C-ATG4B complex and performed molecular dynamics (MD) simulations (up to 1.5 μs). We found that the WT LC3C-ATG4B complex with and without additional LIR interactions between ATG4B and LC3C remained stable. The C-terminal tail of LC3C remained bound and strongly anchored to the active site of ATG4B throughout the simulation. In the complex, the phosphosites S93 and S96 of LC3C (red cartoon in Figure 3F) are in close-proximity to the ATG4B interacting surface (grey surface). LC3C S96 forms a hydrogen-bond interaction with ATG4B E350, and LC3C S93 is close to a network of hydrogen bonds and salt bridges stabilizing. In MD simulations, double phosphorylation of S93 and S96 interfered with these interactions and disrupted the binding interface between ATG4B and LC3C (**Movies SM1, SM2**). The phosphorylated serine residues detached from the ATG4B surface and partially dislodged the LC3C, resulting in partial retraction of the LC3C C-terminal tail from the ATG4B active site. The negative charge introduced by phosphorylation severely weakens complex stability based on calculated binding energies (Table S2), with electrostatic interactions as the dominant factor. Figure S2 shows residue-wise contributions to the binding energy mapped onto the LC3C structure. According to these calculations, phosphorylated S93 and S96 are strongly destabilizing (Figure S2; red thick cartoon), whereas unphosphorylated S93 and S96 are favorable (Figure S2; blue thin cartoon). The MD simulations and binding energy calculations indicate that phosphorylation disrupts the LC3C-ATG4B interface and destabilizes the complex.

### Phosphorylation at S93 and S96 affects LC3C C-terminal tail structure and thereby impedes ATG4-mediated cleavage

Based on the simulation results for the LC3C-ATG4B complex, we hypothesized that phosphorylation of unbound LC3C could affect its C-terminal tail structure and prevent binding to the ATG4B active site. In MD simulations (see Methods) of free LC3C, we found that the C-terminal tail of LC3C (126-147) was disordered and highly dynamic (**Movie SM3**). By contrast, in the phosphorylated variants (S93PO_4_ LC3C and S96-PO_4_ LC3), the C-terminal tail adopted more ordered conformations (**Movies SM4, SM5**; Figure 4A-B). The phosphoserines formed intramolecular salt bridges with R134 (Figures 4A and 4B) that pulled the C-terminal tail of LC3C towards the protein, structuring it locally. In repeated simulations (*n* = 6 each) of unphosphorylated and phosphorylated variants of LC3C (Figures S3A- C), we observed a total of six salt-bridge formation events, indicating that the intramolecular salt-bridge formation between the phosphoserines and R134 is robust. We observed the salt bridge formation on a sub-microsecond time scale (Figure 4D and 4E). To confirm this finding and the role of R134, we performed ATG4mediated *in vitro* cleavage experiments of double tagged LC3C WT, S93/96D, S93/96D R134A, and S93/96D R142A (a control mutation in the C-terminal tail). The LC3C C-terminal cleavage was monitored by the disappearance of its C-terminal Strep-tag. The mutation of S93/96D delayed the cleavage of the C-terminal tail of LC3C by ATG4B (Figure 4F). The R134A mutation could partially rescue this phenotype of S93/96D, whereas the other C-terminal tail mutation, R142A, could not (Figure 4F). The results of the ATG4-mediated cleavage assay are thus consistent with R134-phosphoserine interactions sequestering the LC3C C-terminal tail and preventing access to ATG4B and subsequent cleavage.

**Figure 4:**
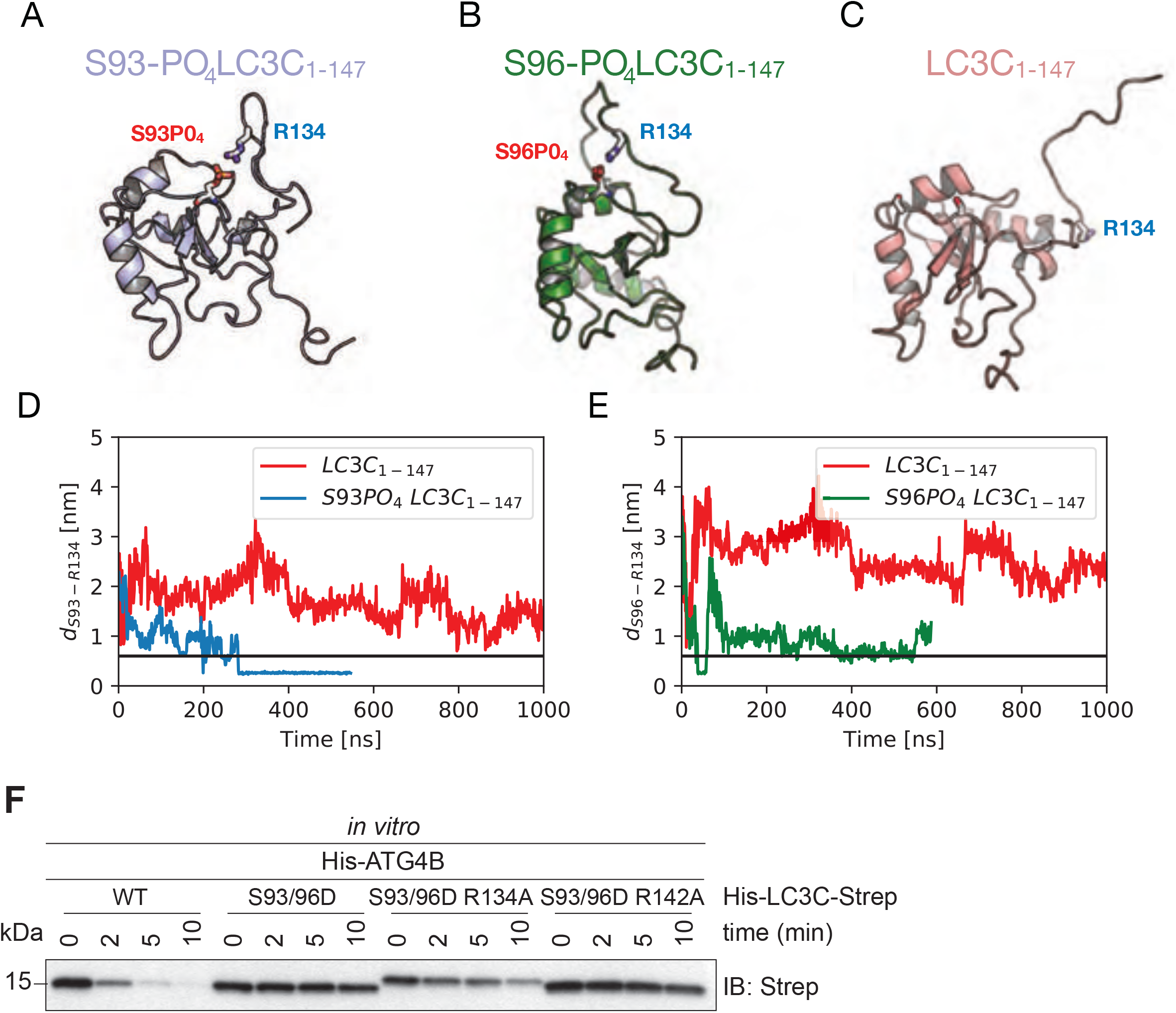
Phosphorylation at S93 and S96 affects LC3C C-terminal tail structure in pro-LC3C, thereby impeding ATG4-mediated processing. (A-C) Representative snapshots from all-atom MD simulations (see Movies SM3-5) of unphosphorylated (red), S93-PO_4_ LC3C (blue) and S96-PO_4_ LC3C (green). (D, E) Salt-bridge formation dynamics in MD simulations represented by the time-dependent minimum distance between side chain heavy atoms of R134 to (D) S93 and (E) S96 in phosphorylated (blue) and unphosphorylated (red) LC3C simulations. Black line represents cut-off distance (0.6 nm) for favorable salt-bridge formation. (F) SDS-PAGE and Western blotting of *in vitro* ATG4 cleavage assay. Purified double tagged His-LC3C-Strep WT and mutants were incubated with ATG4B in buffer for indicated time points. LC3C S93/96D R134A mutation enables C-terminal cleavage of LC3C by ATG4B.

### Phospho-mimetic LC3C and GABARAP-L2 cannot form autophagosomes in cells

GABARAP-L2 lacks the C-terminal tail, and the ATG4B-mediated processing removes only a single C-terminal residue (F117), which exposes G116 for lipidation. Therefore, we hypothesized that phosphorylating S87 and S88 in GABARAP-L2 weakens binding to ATG4B and in turn slows down proteolytic processing. Accordingly, we tested if phospho-mimetic GABARAP-L2 S88D and LC3C S93/96D can form autophagosomes, despite not being processed by ATG4B. U2OS cells were co-transfected with HA-Parkin and GFP-GABARAP-L2 WT, S88D, S88A (Figure 5A and Figure S4A) or GFP-LC3C WT, S93/96D, S93/96A (Figure 5B and Figure S4B) and mitophagy was induced by the addition of CCCP for 3 hours. Upon induction of mitophagy GABARAP-L2 WT and S88A formed autophagosomes. By contrast, the phospho-mimetic GABARAP-L2 S88D remained dispersed throughout the cell and no autophagosome formation was observed (Figure 5A and Figure S4A). Likewise, phospho-mimetic LC3C S93/96D did not form autophagosomes upon induction of mitophagy unlike WT and S93/96A LC3C (Figure 5B and Figure S4B).

**Figure 5:**
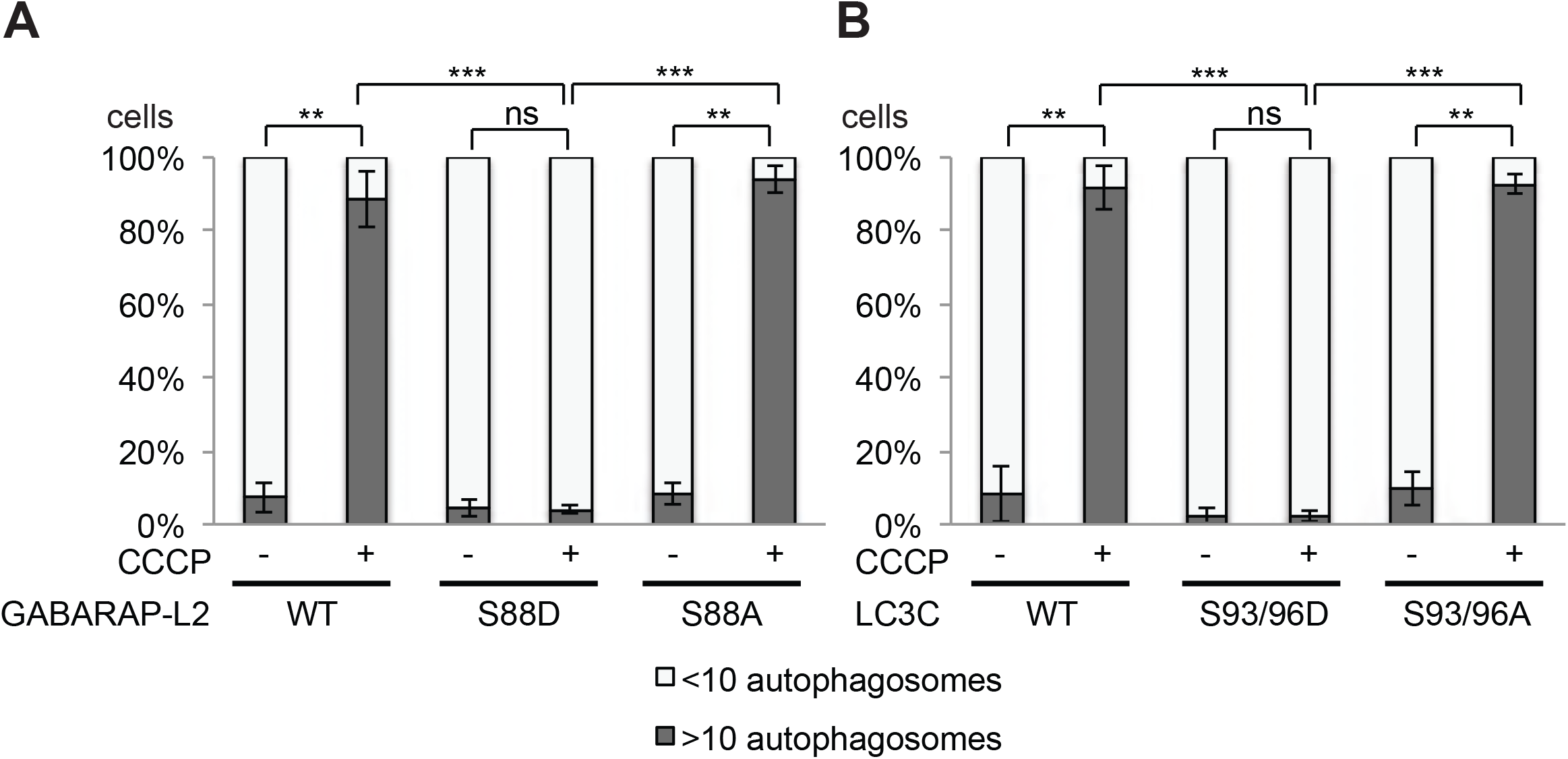
Phospho-mimetic LC3C and GABARAP-L2 cannot form autophagosomes. (A,B) U2OS cells were transfected with GFP-GABARAP-L2 (A) or GFP-LC3C (B) WT or mutants and HA-Parkin. Mitophagy was induced by the addition of 40 μM CCCP for 3 hours. GFP-expressing cells were counted and segregated into classes with greater and less than 10 autophagosomes per cell. Bars represent mean±SD from three separate experiments. * P<0.05, ** P<0.01, ***P<0.001, as analyzed by unpaired Student’s t-test.

### Phospho-mimetic Δ C-terminal LC3C and GABARAP-L2 are not lipidated and do not form autophagosomes

Since LC3 family proteins can only be integrated into autophagosomes after C-terminal cleavage by ATG4, we tested whether artificially truncated LC3C or GABARAP-L2 (Δ C-term: LC3C (1-126) and GABARAP-L2 (1-116)) could circumvent ATG4-mediated processing, undergo lipidation, and form autophagosomes. U2OS cells were co-transfected with HA-Parkin and GFP-GABARAP-L2 Δ C-term WT, S88A or S88D (Figure 6A and Figure S5A) or GFP-LC3C Δ C-term WT, S93/96A or S93/96D (Figure 6B and Figure S5B) and mitophagy was induced by the addition of CCCP for 3 hours. Upon induction of mitophagy, GABARAP-L2 or LC3C Δ C-term WT and alanine mutants formed autophagosomes, while phosphomimetic mutants with truncated C-terminus (GABARAP-L2 Δ C-term S88D or LC3C Δ C-term S93/96D) remained dispersed throughout the cell and no autophagosome formation could be observed (Figure 6A and Figure 6B). Upon induction of mitophagy, GABARAP-L2 lipidation can be observed by the appearance of a lower band on Western Blots (Figure 3D), which can also be observed during mitophagy induction of GABARAP-L2 Δ C-term WT and S88A, but not with phosphomimetic GABARAP-L2 Δ C-term S88D (Figure S5C). Hence, the phosphorylation of LC3C or GABARAP-L2 not only impedes their C-terminal cleavage by ATG4, but also their lipidation by the lipidation cascade enzymes ATG12-5-16L1. ATG7 function is similar to ubiquitin-activating (E1) enzymes; it recruits ATG3 (an E2-like enzyme), which then catalyzes the conjugation to the lipid moiety (PE) to the C-terminal exposed glycine of the truncated LC3 Δ C-term. Binding of the ATG12-5-16L1 complex (E3-like enzyme) to ATG3 enhances the lipidation of LC3, since the ATG5-ATG12 complex ensures that nascently lipidated LC3 is incorporated into the phagophore membrane (Nakatogawa, 2013). In order to test if phosphorylation of LC3C/GABARAP-L2 Δ C-term impedes their processing by the lipidation cascade enzymes, we also performed an *in vitro* lipidation assay (Figure 6C). LC3C Δ C-term WT and LC3C Δ C-term S93/96D or GABARAP-L2 Δ C-term WT and GABARAP-L2 Δ C-term S87/88D were incubated with ATP, liposomes, hATG3, hATG7, and hATG12-5-16L1 (reaction mix). WT LC3C and GABARAP-L2 could be successfully lipidated, while S93/96D LC3C and S87/88D GABARAP-L2 could not be conjugated to membrane *in vitro* (Figure 6C,D) indicating that phosphorylation of LC3s also affects lipid conjugation.

**Figure 6:**
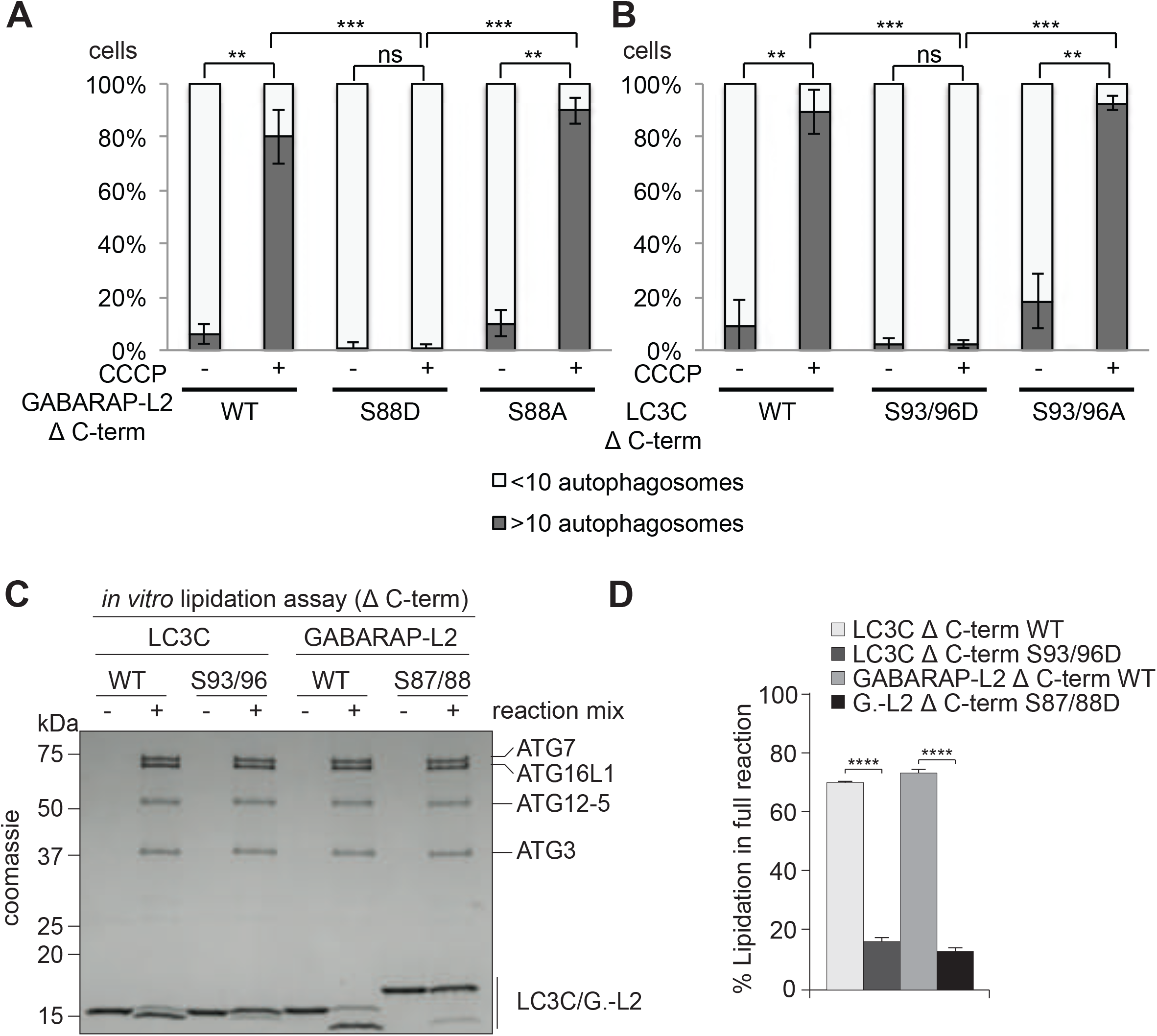
Phospho-mimetic Δ C-terminal LC3C or GABARAP-L2 cannot form autophagosomes because they cannot be lipidated. (A,B) U2OS cells were transfected with GFP-GABARAP-L2 Δ C-terminal (A) or GFP-LC3C Δ C-terminal (B) WT or mutants and HA-Parkin. Mitophagy was induced by the addition of 40 μM CCCP for 3 hours. GFP-expressing cells were counted and segregated into classes with greater and less than 10 autophagosomes per cell. Data is presented as mean±SD from three separate experiments. *P<0.05, **P<0.01, ***P<0.001 as analyzed by Students T-test. (C) *In vitro* lipidation reactions containing 10 μM LC3C WT, LC3C S93/96D, GABARAP-L2 (GL2) WT or GABARAP-L2 S87/88D incubated with or without 0.5 μM hATG7, 1 μM hATG3, 0.25 µM hATG12-5-16L1β, 3 mM lipid (sonicated liposomes composed of 10 mol% bl-PI, 55 mol% DOPE, and 35 mol% POPC), 1 mM DTT and 1 mM ATP were incubated at 30°C for 90 minutes. The reactions were analyzed by SDS-PAGE and visualized by Coomassie blue stain. (D) The extent of lipidation in (C) was quantified and plotted as percentage of total protein (conjugated and unconjugated). Data is presented as mean±SEM from three separate experiments. *P<0.05, **P<0.01, ***P<0.001, ****P<0.0001, as analyzed by One-way Anova followed by Bonferonis multiple comparison test.

### TBK1-mediated GABARAP-L2 phosphorylation impedes its premature cleavage from autophagosomes by ATG4

TBK1 is recruited to the site of autophagosome formation by autophagy receptor proteins, where TBK1 phosphorylates OPTN and p62 to promote autophagy flux (Heo et al., 2015, Lazarou et al., 2015, Matsumoto et al., 2011, Ordureau et al., 2015, Pilli et al., 2012, Richter et al., 2016a, Thurston et al., 2009, Wild et al., 2011). Hence, it is most likely that LC3C and GABARAP-L2 are phosphorylated by TBK1 during autophagosome formation and not during the initial processing step of pro-LC3 cleavage post ribosomal release. In order to test whether the TBK1-mediated phosphorylation of LC3C and GABARAP-L2 has an impact on ATG4-mediated de-lipidation of LC3s from the mature autophagosome, an *in vitro* delipidation assay of LC3C and GABARAP-L2 proteins was performed. The fractions of PE-conjugated LC3C Δ C-term WT, phospho-mimetic S93/96D and PE-conjugated GABARAP-L2 Δ C-term WT and phospho-mimetic S87/88D (see Methods) were enriched and used as substrates for the de-lipidating enzymes ATG4A or ATG4B (Figure 7A,B). We found that neither LC3C Δ C-term WT, nor LC3C Δ C-term S93/96D could be de-lipidated and released from the liposome by ATG4A (Figure 7A,B). In contrast, ATG4B is able to de-lipidate and cleave small amounts of LC3C Δ C-term WT (Figure S6A) from liposomes, but has no activity towards LC3C Δ C-term S93/96D. This indicates that lipidated LC3C could be targeted specifically by other isoforms of ATG4 enzymes (ATG4C or ATG4D) but not by ATG4A and ATG4B. WT GABARAP-L2 could be de-lipidated from liposomes by ATG4B and ATG4A (at a slower rate) in a dose dependent manner (Figure S6B). On the contrary, we found that GABARAP-L2 S87/88D is not a target of ATG4A or B (Figure 7A,B).

**Figure 7:**
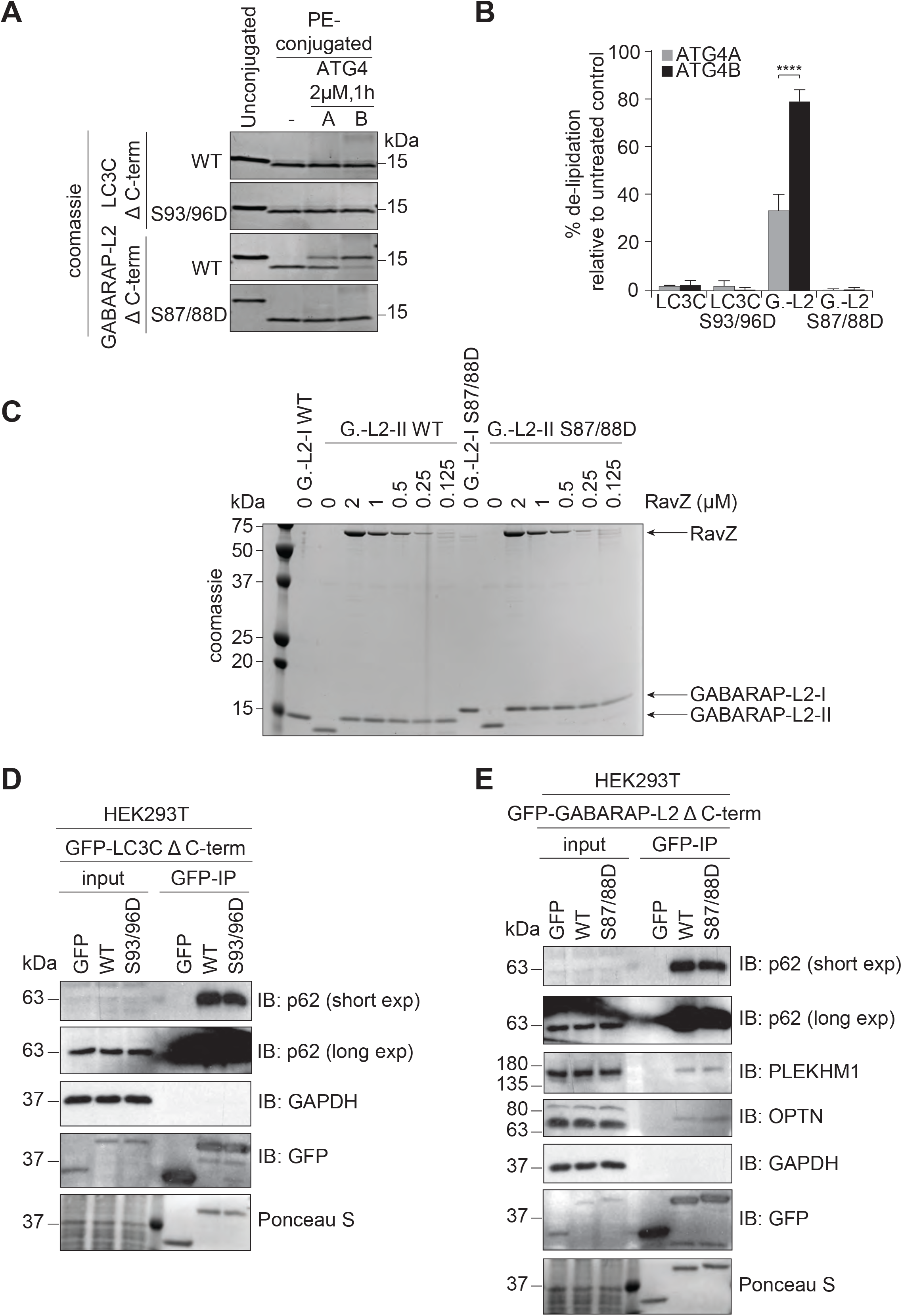
TBK1-mediated GABARAP-L2 phosphorylation impedes its premature cleavage from autophagosomes by ATG4. (A) LC3C WT-, LC3C S93/96D, GABA-RAP-L2 (G.-L2) WT- or GABARAP-L2 S87/88D-conjugated liposomes were treated or not with 2 µM ATG4A or ATG4B for 1 hour at 37°C. Samples were then subjected to SDS-PAGE together with unconjugated LC3C and GABARAP-L2 proteins. (B) The extent of de-lipidation in (A) was quantified and plotted as percentage of total protein (conjugated and unconjugated). Data is presented as mean±SEM from three separate experiments. *P<0.05, **P<0.01, ***P<0.001, ****P<0.0001, as analyzed by Two-way Anova followed by Bonferonis multiple comparison test. (C) GABARAP-L2 WT- or GABARAP-L2 S87/88D-conjugated liposomes (G.-L2-II) were treated or not with different amounts (2-0.125 µM) of RavZ for 1 hour at 37°C. Samples were then subjected to SDS-PAGE together with un-conjugated GABARAP-L2 (G.-L2-I). (D) SDS-PAGE and Western blot of HEK293T cell lysates and GFP IPs. Cells were transfected with GFP, GFP-LC3C Δ C-terminal WT or S93/96D and lysates used for GFP IPs. WT and S93/96D LC3C bind endogenous p62. (E) SDS-PAGE and Western blot of HEK293T cell lysates and GFP IPs. Cells were transfected with GFP, GFP-GABARAP-L2 Δ C-terminal WT or S87/88D and lysates used for GFP IPs. WT and S87/88D GABARAP-L2 bind endogenous p62, Optineurin (OPTN) and PLEKHM1.

In addition, we also tested whether the phosphorylation of GABARAP-L2 has an impact on its de-lipidation from the phagophore by other proteases such as RavZ (Figure 7C). RavZ is a bacterial effector protein from the intracellular pathogen *Legionella pneumophila* that interferes with autophagy by directly and irreversibly uncoupling GARARAP-L2 attached to PE on autophagosome membranes (Choy et al., 2012, Kwon et al., 2017). We found that small amounts of RavZ could remove GARABAP-L2 WT and S87/88D mutant from autophagosomes (Figure 7C), indicating its effectiveness in circumventing *Legionella* growth restriction via xenophagy (when TBK1 is also activated). Likewise, RavZ is also able to cleave LC3C WT and S93/96D mutant from liposomes *in vitro* (Figure S6A).

Finally, we tested if phosphorylated LC3C or GABARAP-L2 adhered to autophagosomes are still functional to perform downstream reactions. LC3 family proteins interact with autophagosome receptors such as p62, which link the growing autophagosome to cargo (Pankiv et al., 2007, Zheng et al., 2009). Both LC3C Δ C-term WT and phospho-mimetic S93/96D can bind to p62 (Figure 7D). Similarly, p62 and OPTN can be recruited to autophagosomes by WT as well as S87/88D GABARAP-L2 (Figure 7E) (Wild et al., 2011, Wong & Holzbaur, 2014). Once all of the cargo has been engulfed by the autophagosome, degradation can take place through the fusion with lysosomes (Nakamura & Yoshimori, 2017). GABARAP family proteins mediate autophagosomal-lysosomal fusion by binding to the autophagy adaptor protein PLEKHM1 (McEwan et al., 2015, Wang et al., 2015). Phosphorylation of GABARAP-L2 by TBK1 does not interfere with its ability to bind to PLEKHM1 (Figure 7E).

Hence, TBK1 mediated phosphorylation of GABARAP-L2 and LC3C protects them from premature autophagosome removal by ATG4, but does not interfere with downstream reactions like cargo binding and lysosomal fusion.

## Discussion

The autophagy pathway is tightly regulated to ensure proper recycling and disposal of cellular material during nutrient shortage. Here, we present a new regulatory mechanism of autophagy, which influences the conjugation and de-conjugation of LC3C and GABARAP-L2 to autophagosomes. The kinase TBK1 fulfils several roles during selective autophagy. Upon autophagy induction, TBK1 is recruited to the site of autophagosome formation and gets activated by trans-autophosphorylation after accumulation (Ma et al., 2012, Shu et al., 2013). We show that, at this stage, TBK1 can phosphorylate LC3C and GABARAP-L2 at specific serine residues to protect them from ATG4-mediated premature removal from autophagosomes.

ATG4 mediates regular processing of pro-LC3s post ribosomal release, de-lipidation of incorrectly lipidated LC3-PE on other endomembranes, and favors incorporation of LC3s into autophagosomes by ATG12-5-16L1. Spatial and temporal regulation of recruitment and dissociation of LC3 family proteins to and from autophagosomes is achieved through regulation of ATG4 activity (Pengo et al., 2017, Sanchez-Wandelmer et al., 2017, Scherz-Shouval et al., 2007, Yang et al., 2015, Yu et al., 2012). ATG4 constitutively de-conjugates LC3 family proteins from all endomembranes except from autophagosomes, to maintain a pool of unlipidated LC3 (Nakatogawa et al., 2012). This suggests that LC3-PE conjugated to autophagosomes is protected from premature de-lipidation by a timely regulatory mechanism. This regulation is achieved through the phosphorylation and dephosphorylation of ATG4 itself (Pengo et al., 2017, Sanchez-Wandelmer et al., 2017) and the phosphorylation of LC3s by TBK1.

The kinase activity of TBK1 is tightly regulated (Xu 2017) and during xenophagy and mitophagy, TBK1 phosphorylates autophagy receptor proteins (Heo et al., 2015, Lazarou et al., 2015, Ordureau et al., 2015, Pilli et al., 2012, Richter et al., 2016a, Thurston et al., 2009, Wild et al., 2011).

The active recruitment of TBK1 to the sites of autophagosome formation (Lazarou et al., 2015, Richter et al., 2016a) makes it likely that TBK1-mediated phosphorylation occurs on nascent phagophores, resulting in phosphorylated forms of membrane-embedded LC3s. Phosphorylation prevents premature removal of lipidated LC3C/GABARAP-L2 from growing autophagosomes by ATG4. Molecular modeling and atomistic simulations of the ATG4B-LC3C complex revealed that LC3C phosphorylation impedes binding to ATG4. The weakened binding slows down delipidation, which ensures that a steady coat of lipidated LC3C/GABARAP-L2 is maintained throughout the early steps in autophagosome formation (Figure 8). The phosphorylation of LC3C/GABARAP-L2 does not impede their binding to autophagy receptors such as p62 or PLEKHM1, which promotes unhindered downstream steps for, e.g., autophagosome-lysosome fusion (McEwan et al., 2015).

**Figure 8:**
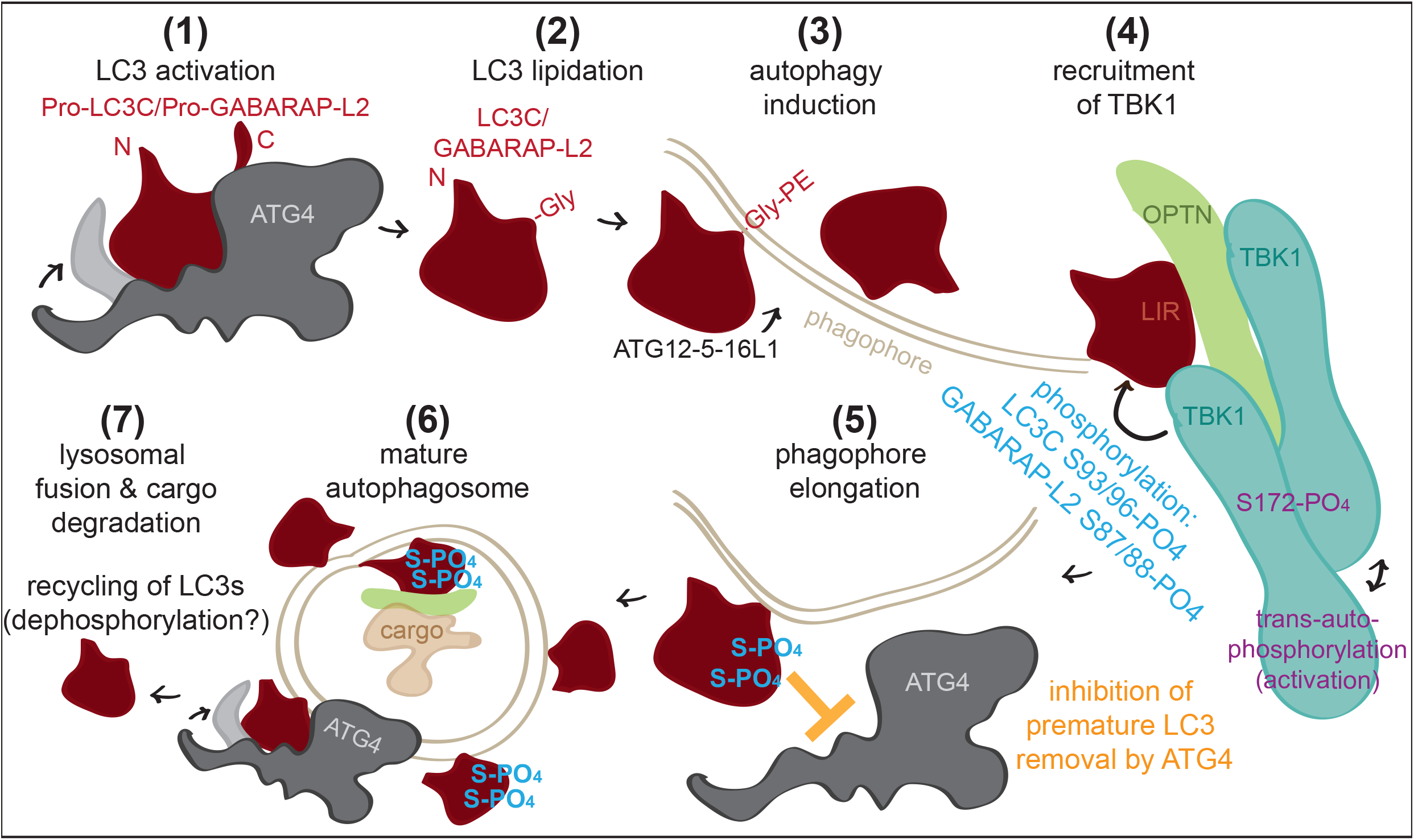
Model of TBK1-mediated LC3C and GABARAP-L2 phosphorylation. TBK1 recruitment to autophagosomes promotes phosphorylation of membrane-embedded LC3C and GABARAP-L2. Phosphorylation prevents premature removal of LC3C and GABARAP-L2 from autophagosomes, ensuring an unperturbed and unidirectional flow of the autophagosome to the lysosome.

Thus, phosphorylation of LC3s aids in maintaining an unperturbed and unidirectional flow of the autophagosome to the lysosome.

At later stages of autophagosome formation, this process could be slowed-down or reversed by either TBK1 dissociation from autophagosomes or diminished catalytic activity. Alternatively, action of phosphatases could allow de-lipidation prior to autophagosomal-lysosomal fusion, thereby recycling LC3s.

## Materials and Methods

### Expression constructs

Expression constructs of indicated proteins were cloned into indicated vectors using PCR or the gateway system. Site-directed mutagenesis was performed by PCR to introduce desired amino acid substitutions. All expression constructs were sequenced by Seqlab.

### Protein expression and purification

GST or His-tagged fusion proteins were expressed in *E. coli* strain BL21 (DE3). Bacteria were cultured in LB medium supplemented with 100 μg/mL ampicillin at 37°C in a shaking incubator (150 rpm) until OD600 ∼0.5-0.6. Protein expression was induced by the addition of 0.5 mM IPTG and cells were incubated at 16°C for 16 hours. Bacteria were harvested by centrifugation (4000 rpm) and lysed by sonication in GST lysis buffer (20 mM Tris HCl, pH 7.5, 10 mM EDTA, pH 8.0, 5 mM EGTA, 150 mM NaCl, 0.1% β-mercaptoethanol, 1 mM PMSF) or His lysis buffer (25 mM Tris HCl, pH 7.5, 200 mM NaCl, 0.1% β-mercaptoethanol, 1 mM PMSF, 1mg/ml lysozyme). For the purification of ATG4 the use of PMSF was omitted. Lysates were cleared by centrifugation (10000 rpm), 0.05% of Triton X-100 was added and the lysates were incubated with glutathione Sepharose 4B beads (GE Life Sciences) or Ni-NTA agarose beads (Thermo Fisher) on a rotating platform at 4°C for 1 hour. The beads were washed five times either in GST wash buffer (20 mM Tris HCl, pH 7.5, 10 mM EDTA, pH 8.0, 150 mM NaCl, 0.5% Triton X-100, 0.1% β-mercaptoethanol, 1 mM PMSF) or His wash buffer (25 mM Tris HCl, pH 7.5, 200 mM NaCl, 0.05% Triton X-100, 10 mM Imidazole). The immobilized proteins were reconstituted in GST storage buffer (20 mM Tris HCl, pH 7.5, 0.1% NaN3, 0.1% β-mercaptoethanol) or eluted with His elution buffer (25 mM Tris HCl, pH 7.5, 200 mM NaCl, 300 mM Imidazole) and dialysed in (25 mM Tris HCl, pH 7.5, 200 mM NaCl) at 4°C over night.

Recombinant GST-TBK1 was obtained from the MRC PPU DSTT in Dundee, UK (#DU12469).

Purification of proteins used for *in vitro* lipidation/de-lipidation: Full-length hATG7, hATG3 and hATG5-12–ATG16L1 complex was expressed and purified from HEK suspension cells (HEK-F, Invitrogen) as previously described (Lystad et al., 2019). To purify mATG3, LC3C, LC3C S93/96D, GABARAP-L2 WT, GABARAP-L2 S87/88D and RavZ, pGEX-6P-1 plasmid containing the corresponding cDNA was transformed into BL21-Gold (DE3) E. coli. Cells were grown at 37°C to an OD of 0.6-0.8 before induction with 0.5 mM IPTG. Cells were then grown for 3 additional hours before they were collected by centrifugation. Cells were resuspended in NT350 (20 mM Tris-HCl pH 7.4, 350 mM NaCl) supplemented with a Roche Complete Protease inhibitor, lysed by sonication and cleared by centrifugation. The supernatant was incubated at 4ºC with Glutathione Beads (Sigma) for 4 hours. Beads were collected and washed twice with NT350 buffer before HRV 3C protease was added and allowed to cut at 4°C overnight. The next morning, protein fractions were collected and stored at −80°C in 20% glycerol. (mATG3 and RavZ plasmids were a gift from Thomas Melia (Yale University)).

### Cell culture

HEK293T and U2OS cells were cultured in Dulbecco’s modified Eagle’s medium (DMEM; Gibco) supplemented with 10% fetal bovine serum, 2 mM L-glutamine, and 1% penicillin/streptomycin and maintained at 37°C in a humidified atmosphere with 5% CO_2_. CCCP was resuspended in DMSO and cells were treated with 40 μM for 3 hours. Plasmid transfections were performed with 3 μl GeneJuice (Merck Millipore), 0.5 μg plasmid DNA in 200 μl Opti-MEM (Life Technologies). After incubation for 15 min, the solution was added to the cells, which were lysed in lysis buffer or fixed with 4% paraformaldehyde 48 hours later.

### Immunofluorescence microscopy

Transfected U2OS cells were seeded onto glass coverslips in 12-well culture dishes and treated accordingly. Cells were washed in phosphate-buffered saline (PBS) before fixation with 4% paraformaldehyde for 10 minutes at room temperature. The coverslips were washed a further three times before permeabilization of the cells with 0.5% Triton X-100 in PBS for 10 minutes at room temperature. Cells were rinsed with PBS before being incubated for 1 hour in 1% bovine serum albumin (BSA) in PBS for 1 hour. Primary antibody incubation was done for 1 hour in a humidified chamber with 1% BSA in PBS. After thorough washes in PBS, cells were incubated with Alexa Fluor secondary antibodies 1% BSA in PBS for 1 hour in the dark. Cells were washed three more times in PBS and once with deionized water before being mounted onto glass slides using ProLong Gold mounting reagent (Life Technologies), which contained the nuclear stain 4′,6-diamidino-2-phenylindole (DAPI). Slides were imaged using a Leica microscope Confocal SP 80 fitted with a 60× oil-immersion lens.

### Cell lysis

For lysis, cells were scraped on ice in lysis buffer (50 mM Hepes, pH 7.5, 150 mM NaCl, 1 mM EDTA, 1 mM EGTA, 1% Triton X-100, 25 mM NaF, 5% glycerol, 10μM ZnCl_2_) supplemented with complete protease inhibitors (cOmplete, EDTA-free; Roche Diagnostics) and phosphatase inhibitors (P5726, P0044; Sigma-Aldrich). Extracts were cleared by centrifugation at 15000 rpm for 15 minutes at 4°C.

### Immunoprecipitation and protein binding assays

Cleared cell extracts were mixed with glutathione-Sepharose beads (GE Healthcare) conjugated to LC3 family proteins or GST, FLAG-agarose beads (Sigma-Aldrich) or GFP-Trap_A beads (ChromoTek) for 2 hours at 4°C on a rotating platform. The beads were washed four times in lysis buffer. Immunoprecipitated and input samples were reduced in SDS sample buffer (50 mM Tris HCl, pH 6.8, 10% glycerol, 2% SDS, 0.02% bromophenol blue, 5% β-mercaptoethanol) and heated at 95°C for 5 minutes (Herhaus et al., 2013, Herhaus et al., 2014).

### Western blotting

For immunoblotting, proteins were resolved by SDS-PAGE and transferred to PVDF membranes. Blocking and primary antibody incubations were carried out in 5% BSA in TBS-T (150 mM NaCl, 20 mM Tris, pH 8.0, 0.1% Tween-20), secondary antibody incubations were carried out in 5% low-fat milk in TBS-T and washings in TBS-T. Blots were developed using Western Blotting Luminol Reagent (sc-2048; Santa Cruz). All Western blots shown are representative.

### Antibodies

The following antibodies were used in this study: anti-HA-tag (11867423001; Roche), anti-FlagM2-tag (F3165; Sigma-Aldrich), anti-GFP-tag (Living Colors 632592; Clontech), anti-Strep-tag (34850; Qiagen), anti-His-tag (11922416001; Roche), anti-vinculin (V4505; Sigma), anti-tubulin (T9026; Sigma), anti-TBK1 (#3013; Cell Signaling Technology), anti-pTBK1 (pS172; #5483; Cell Signaling Technology). Secondary HRP conjugated antibodies goat anti-mouse (sc-2031; Santa Cruz), goat anti-rabbit (sc-2030; Santa Cruz) and goat anti-rat (sc-2006; Santa Cruz), IgGs were used for immunoblotting. Donkey anti-rat Alexa Fluor 647 (A-21247; Life Technologies) was used for immunofluorescence studies.

### Kinase assays

LC3 family proteins were incubated in 20 μL phosphorylation buffer (50 mM Tris HCl, pH 7.5, 10 mM MgCl_2_, 0.1 mM EGTA, 20 mM ß-glycerophosphate, 1 mM DTT, 0.1 mM Na_3_VO_4_, 0.1 mM ATP or γ^32P^ ATP (500 cpm/pmol)) with 50 ng of recombinant GST-TBK1 for 15 minutes at 30°C. The kinase assay was stopped by adding SDS sample buffer containing 1% β-mercaptoethanol and heating at 95°C for 5 minutes. The samples were resolved by SDS-PAGE, and the gels were stained with InstantBlue (Expedeon) and dried. The radioactivity was analysed by autoradiography (Herhaus et al., 2015).

### SILAC-IP and phosphopeptide identification

Cells were maintained in custom-made SILAC DMEM (heavy: R10, K8/light R0, K0) for 14 days, treated accordingly and lysed (as stated above). Incorporation of labeled amino acids to more than 95% was verified by Mass spectrometry. Lysates of SILAC-labeled cells expressing GFP-tagged LC3C or GFP-tagged GABARAP-L2, TBK1 WT (heavy labeled) and TBK1 K38A (light labeled) were combined at equal amounts and incubated with GFP-Trap beads for 1 hour, followed by washes under denaturing conditions (8 M Urea, 1% SDS in PBS). Bound proteins were eluted in NuPAGE LDS Sample Buffer (Life Technologies) supplemented with 1 mM DTT, boiled at 70°C for 10 minutes, alkylated and loaded onto 4-20% gradient SDS-PAGE gels. Alternatively, *in vitro* phosphorylated LC3 family proteins (as stated above) were used to determine TBK1-dependent phosphorylation sites. Proteins were stained using InstantBlue and digested in-gel with trypsin. Peptides were extracted from the gel, desalted on reversed phase C18 StageTips and analyzed on an Orbitrap Elite^TM^ mass spectrometer (ThermoFisher) (Richter et al., 2016a).

### Phos-tag^TM^ SDS-PAGE

Phos-tag^TM^ acrylamide (Wako) gels were used as indicated by the supplier. Gels were prepared with 10% acrylamide, 50 μM phos-tagTM and 100 μM MnCl_2_. Cells were lysed in SDS sample buffer supplemented with 10 μM MnCl_2_.

### ATG4 cleavage assay

Proteins were purified as described above and incubated in buffer (50 mM Tris HCl, pH 7.5, 1 mM EDTA, 150 mM NaCl, 1.5 mM DTT) at 37°C for indicated time points. The assay was stopped by adding SDS sample buffer containing 1% β-mercaptoethanol and heating at 95°C for 5 minutes. The samples were resolved by SDS-PAGE and imaged by Western blotting.

### Modeling and MD simulations with analysis

Starting from the human LC3C (8-125) structure (PDB Id: 3WAM; (Suzuki et al., 2014)), we added N- and C-terminal overhangs using Modeller (Sali & Blundell, 1993) to construct full-length LC3C (1-147). The LC3C-ATG4B complex was modeled using the core complex structure of LC3B(1-124)-ATG4B(1-357) (PDB id: 2Z0E; (Satoo et al., 2009)) as template using Modeller (Sali & Blundell, 1993). The C-terminal tails of both LC3C and ATG4B were modeled in extended conformations without steric clashes across their interface. Additional unresolved loops in ATG4B were modeled using the loop modeling protocol of Modeller (Sali & Blundell, 1993). ATG4B contains an N- and a C-terminal LIR motif, both of which can, in principle, interact with the WXXL-binding sites on either non-substrate or substrate LC3. Therefore, we modeled an alternative WT complex structure including the interaction between the C-terminal LIR of ATG4B and the WXXL-binding site on substrate LC3C. Phosphoserines at S93 and S96 were modeled using CHARMM:GUI (Jo et al., 2008). All structures were solvated in TIP3P water and 150 mM NaCl. After energy minimization, MD simulations of different phosphorylation states were performed using gromacs v5.1 (Pronk et al., 2013), with position-restrained NVT equilibration and NPT equilibration runs for 1000 ps each. Production runs at 310 K and 1 atm were simulated for different times (see Table S1). We used the CHARMM36m force field (Huang et al., 2017), the Nosé-Hoover thermostat (Nosé, 1984), the Parinello-Rahman barostat (Parrinello & Rahman, 1981), and a time step of 2 fs. For each of the LC3C-systems (Table S1), six replicates were simulated with different initial velocities. We also used the molecular mechanics Poisson-Boltzmann surface area (MM-PBSA) to compute the binding energies of the phosphorylated and unphosphorylated LC3C-ATG4B complexes as implemented in g_mmpbsa (Kumari et al., 2014). These binding energies contain molecular mechanical (MM), polar, and non-polar solvation energies. MM energies depend on bonded and non-bonded terms including electrostatic (E_elec_) and van der Waals (E_vdW_) contributions. The polar solvation energies were computed at an ionic strength of 150 mM, a solvent dielectric constant of 80, and a protein dielectric constant of 2 by solving the linearized Poisson-Boltzmann equation with a fine-grid width of 0.5 Å and a coarse grid width of 1.5 times the long axis of the complex, as implemented in Assisted Poisson-Boltzmann Solver (APBS) (Konecny et al., 2012). The non-polar solvation contributions were estimated with the SASA model using a probe radius of 1.4 Å, a surface tension of *γ* = 0.0226 kJ/mol/Å^2^ and an offset of 3.84 kJ/mol (Lee et al., 2000). Binding free energies were estimated as the difference energies between bound and free states, *ΔG*_*Binding*_ = *G*_*LC3C-ATG4B*_ − *G*_*LC3C*_ − *G*_*ATG4B*_, where the free energy contributions of the protein-complex and free proteins are decomposed into a sum of molecular-mechanics, solvent, and configurational entropy contributions,

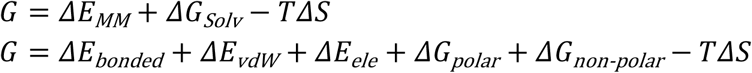

The binding energies were evaluated at intervals of 10 ns from the 1000-ns MD trajectories and averaged (see Table S2). Double differences between unphosphorylated and phosphorylated complexes minimize systematic errors caused by possible energy-function inaccuracy. For the dynamic LC3C-ATG4B protein complexes studied here, these calculated free energy differences point to trends, but should not be interpreted in terms of, say, dissociation constants.

### Liposome and proteoliposome preparation

All lipids were purchased and dissolved in chloroform from Avanti Polar Lipids (Alabaster, AL). Liposomes were prepared by combining 55 mol % 1,2-dioleoyl-sn-glycero-3-phosphoethanolamine (DOPE), 35 mol % 1-palmitoyl-2-oleoyl-sn-glycero-3-phosphocholine (POPC), and 10 mol % bovine liver phosphoinositol (PI). The lipids were dried under nitrogen gas and the lipid film was further dried under vacuum for 1 hour. The lipids were reconstituted in NT350 buffer (350 mM NaCl, 20 mM Tris-HCl pH 7.4) and subjected to 7 cycles of flash-freezing in liquid nitrogen and thawing in a 37°C bath. Liposomes were further sonicated immediately prior to the lipidation reaction.

### Lipidation reaction of LC3C and GABARAP-L2

For a full lipidation reaction LC3C, LC3C S93/96D, GABARAP-L2 WT or GABA-RAP-L2 S87/88D (10 µM) were mixed with hATG7 (0.5 µM), hATG3 (1 µM), Atg12-5-16L1 (0.25 µM), sonicated liposomes (3 mM) and 1 mM DTT. Lipidation was initiated by adding 1 mM ATP and reactions were incubated at 30°C for 90 minutes. Samples were mixed with LDS loading buffer and immediately boiled to stop further lipidation. The reactions were run on a 4-20% SDS-PAGE gel and visualized by Coomassie blue stain and analyzed with Image Lab 6.0 (Biorad).

### De-lipidation reaction of LC3C and GABARAP-L2

For a full lipidation reaction LC3C, LC3C S93/96D, GABARAP-L2 WT or GABA-RAP-L2 S87/88D (10 µM) were mixed with hATG7 (0.5 µM), mAtg3 (containing an extended N-terminal amphipathic helix that permits lipidation in absence of ATG12-5‒16L1) (1 µM), sonicated liposomes (3 mM) and 1 mM DTT. Lipidation was initiated by adding 1 mM ATP and reactions were incubated at 30°C for 90 minutes. After the reaction was complete, the lipidation reaction was run on a Nycodenz density gradient. The bottom layer of the gradient consisted of 150 µL of 80% Nycodenz and 150 µL of the lipidation reaction. The second layer consisted of 250 µL of 30% Nycodenz while the top layer was 50 µL of NT350 buffer. Gradients were centrifuged at 27000 rpm at 4°C for 4 hours in a Beckman SW55 rotor. Liposomes with the conjugated LC3C or GABARAP-L2 protein were collected from the top of the tube before use in subsequent de-lipidation experiments. To measure the activity of proteases, 10 µM of proteoliposomes (concentration estimated by Coomassie blue stain) were mixed with NT350 buffer and kept on ice until activity assays were initiated by the addition of 2 µM (or indicated amounts) of either ATG4A, ATG4B or RavZ. Reactions were incubated at 37°C for 1 hour. Samples were mixed with LDS loading buffer and immediately boiled to stop proteolysis. The reactions were run on a 4-20% SDS-PAGE gel and visualized by Coomassie blue stain and analyzed with Image Lab 6.0 (Biorad).

## Abbreviations

(LIR): LC3 interacting motif
(PE): Phosphatidylethanolamine
(UBD): Ubiquitin-binding domain
(LC3C): MAP1LC3C
(GABARAP-L2): GABA Type A Receptor Associated Protein Like 2
(GATE-16): Golgi-Associated ATPase Enhancer Of 16 KDa
(ATG3): Autophagy Related 3
(ATG5): Autophagy Related 5
(ATG7): Autophagy Related 7
(ATG12): Autophagy Related 12
(ATG16L1): Autophagy Related 16 Like 1
(TBK1): TANK binding kinase 1
(OPTN): Optineurin
(p62): Sequestosome 1
(SDS-PAGE): SDS-polyacrylamide gel electrophoresis
(IP): Immunoprecipitation
(DMEM): Dulbecco’s modified Eagle’s medium
(PBS): Phosphate-buffered saline
(BSA): Bovine serum albumin
(DAPI): 4′,6-diamidino-2-phenylindole
(SILAC): Stable isotope labeling with amino acids in cell culture
(MD): Molecular dynamics
(S-PO_4_): phosphorylated serine

## Acknowledgments

We thank Dr. Masato Akutsu and David McEwan for very valuable comments. This work was supported by grants from the DFG (SFB 1177 on selective autophagy), the Cluster of Excellence "Macromolecular Complexes" of the Goethe University Frankfurt (EXC 115), LOEWE grant Ub-Net and LOEWE Centrum for Gene and Cell Therapy Frankfurt. L.H. is supported by a European Molecular Biology Organization (EMBO) long-term postdoctoral fellowship (ALTF 1200-2014, LTFCO-FUND2013, GA-2013-609409). R.M.B. and G.H. acknowledge support by the Max-Planck Society and computational resources at MPCDF, Garching. A.H.L. and A.S. were supported by the Research Council of Norway (project number 221831) and through its Centers of Excellence funding scheme (project number 262652), as well as the Norwegian Cancer Society (project number 171318).

## Author contributions

LH and ID conceived the study. LH designed and performed most of the experiments. RMB developed structural models and performed MD simulations and analysis of the data with help and supervision from GH. AHL performed *in vitro* (delipidation assays in the lab of AS. LH wrote the manuscript with contribution from all authors. All authors approved the final version of the manuscript.

## Supplementary Figure legends

**Supplementary Figure 1:**
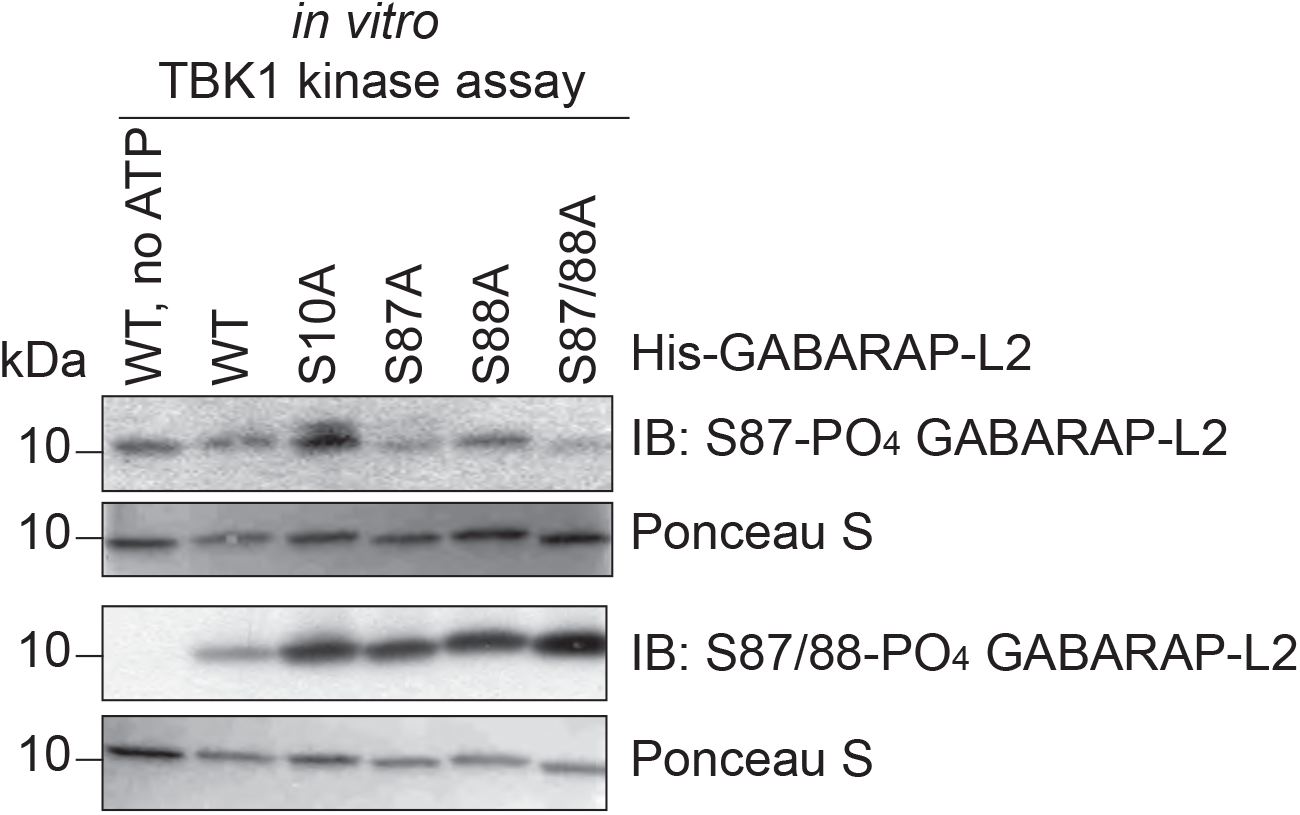
phospho-GABARAP-L2 antibody validation. SDS-PAGE and Western blot of *in vitro* TBK1 kinase assay with His-GABARAP-L2 WT or mutants as substrates to test the respective phospho-GABARAP-L2 antibodies for their specificity.

**Supplementary Figure 2:**
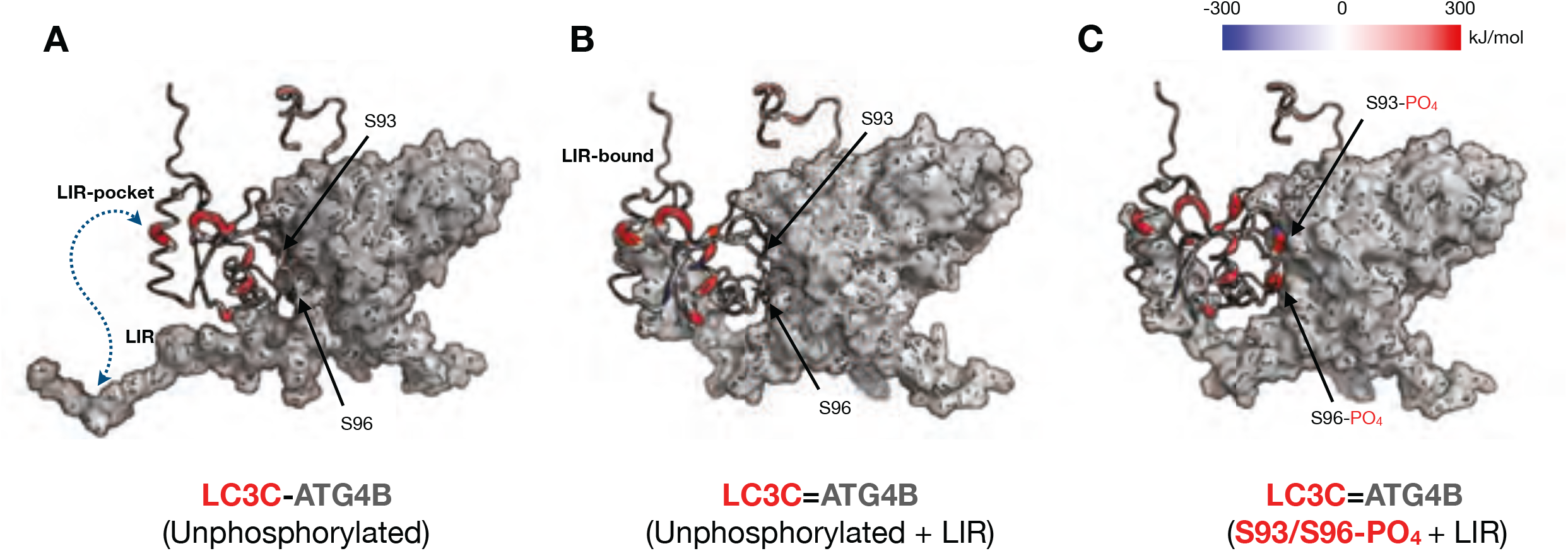
Phosphorylation of S93 and S96 of LC3C affects ATGB binding energy. (A) WT LC3C-ATG4B complex (B) with additional LIR interactions and (C) with phosphorylated LC3C residues (S93 and S96) subjected to MD simulations and binding free energy computations using MM-PBSA (see Methods) approach. A residuewise decomposition of the total binding free energy mapped onto the LC3C structure displays locally favorable (blue), neutral (white) and unfavorable (red) residue interaction with ATG4B (grey surface). Note that S93 and S96 positions in WT complexes contribute favorably (blue color), whereas in the phosphorylated complex contribute unfavorably (red) towards complex formation. The thickness of the backbone scales linearly with the binding energy of LC3C-ATG4B complexes.

**Supplementary Figure 3:**
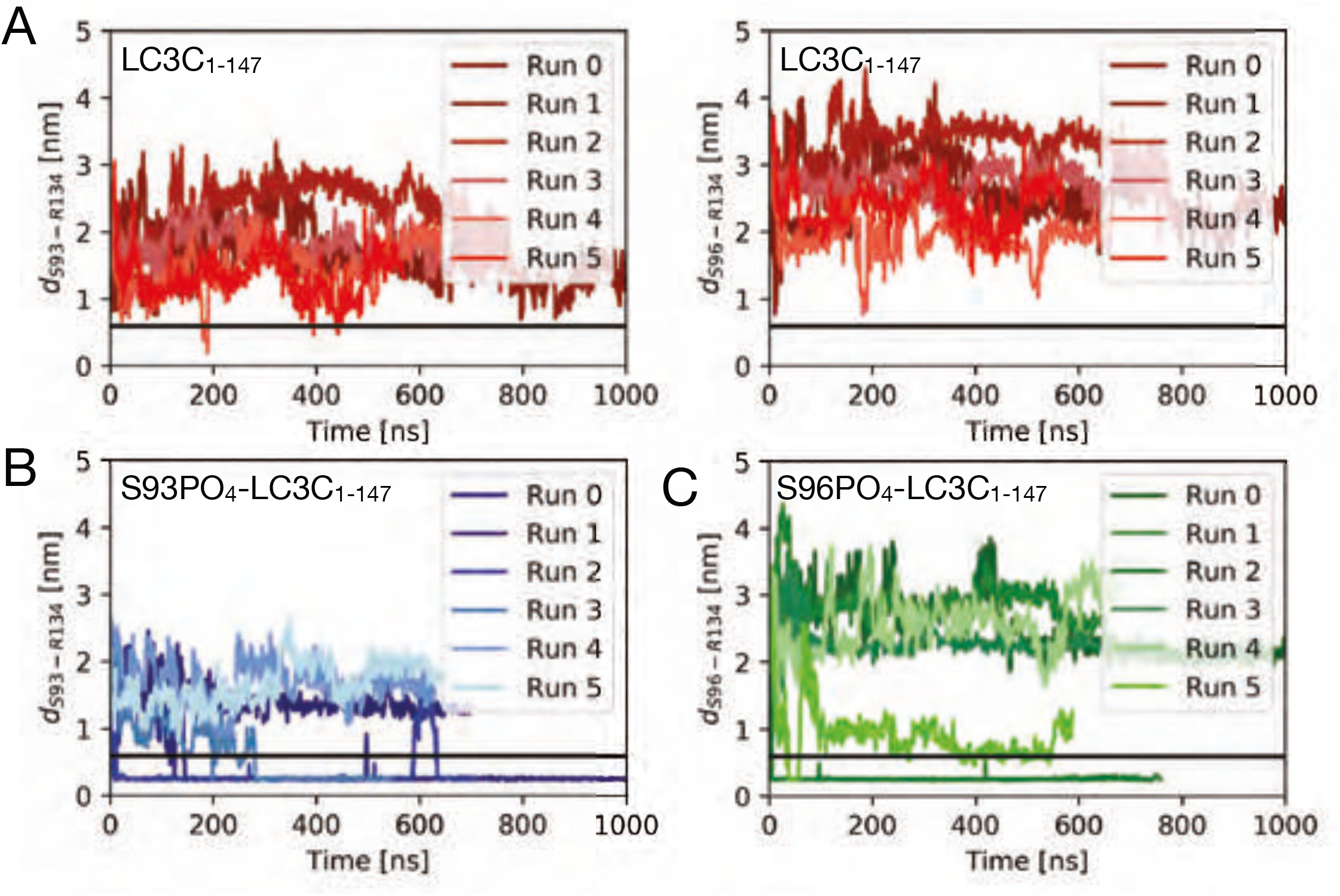
Phosphorylation at S93 and S96 affects LC3C C-terminal tail structure, thereby impeding ATG4-mediated cleavage. Plots of the minimum distance between R134 and serines S93 and S96 report on salt-bridge formation in MD simulations of the LC3C-ATG4B complex with (A) unphosphorylated S93 and S96, (B) S93-PO_4_, and (C) S96-PO_4_. N=6.

**Supplementary Figure 4:**
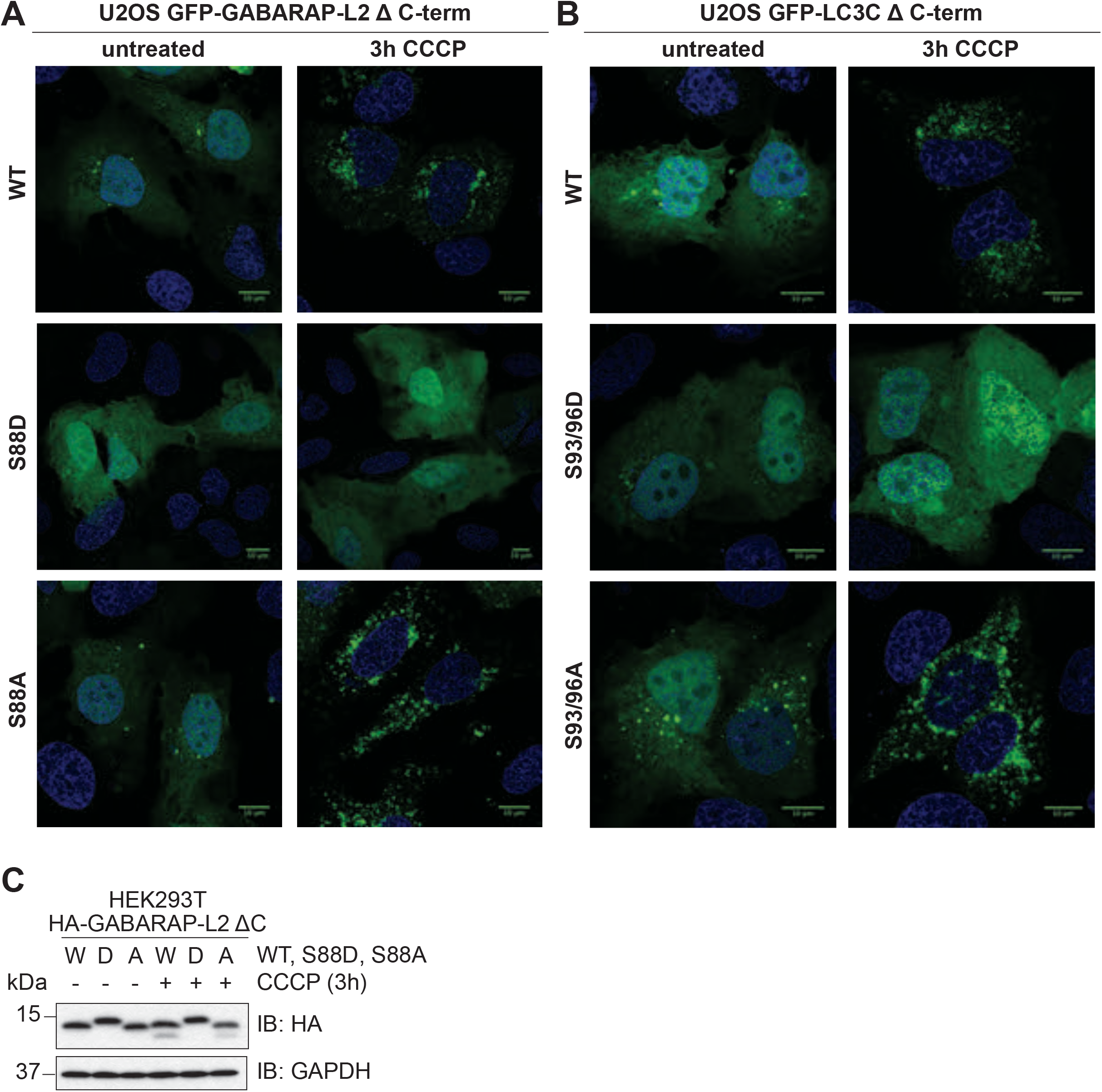
Phospho-mimetic LC3C and GABARAP-L2 cannot form autophagosomes. (A,B) U2OS cells were transfected with GFP-GABARAP-L2 (A) or GFP-LC3C (B) WT or mutants and HA-Parkin. Mitophagy was induced by the addition of 40 μM CCCP for 3 hours. WT and S87/88A GABARAP-L2 form autophagosomes, whereas S87/88D GABARAP-L2 remains dispersed in the cytosol (A). WT and S93/96A LC3C form autophagosomes that localize with HA-Parkin, whereas S93/96D LC3C remains dispersed in the cytosol (B).

**Supplementary Figure 5:**
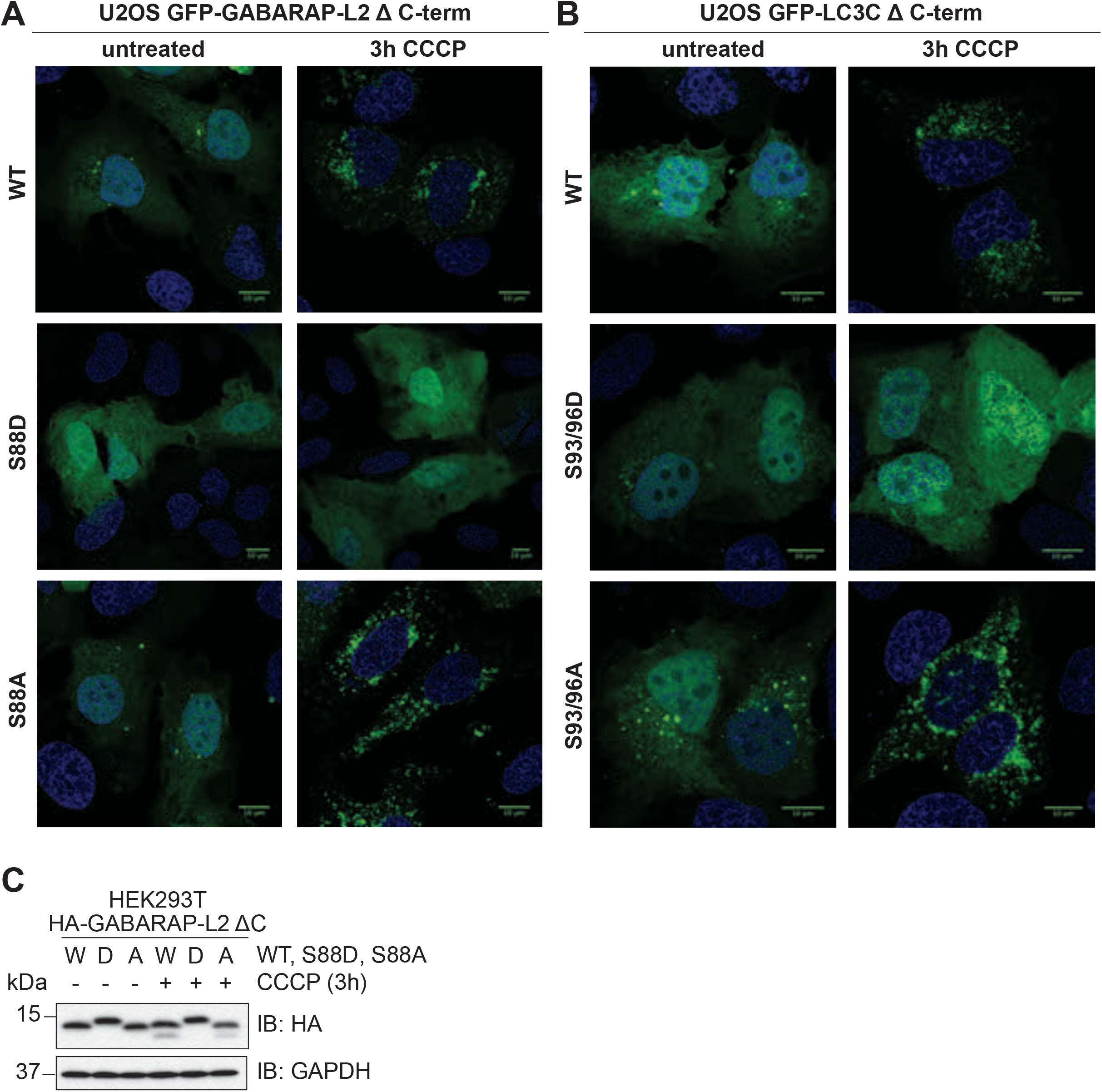
Phospho-mimetic Δ C-terminal LC3C or GABARAP-L2 cannot form autophagosomes. (A,B) U2OS cells were transfected with GFP-GABARAP-L2 Δ C-terminal (A) or GFP-LC3C Δ C-terminal (B) WT or mutants and HA-Parkin. Mitophagy was induced by the addition of 40 μM CCCP for 3 hours. WT and S87/88A Δ C-terminal GABARAP-L2 form autophagosomes, whereas S87/88D Δ C-terminal GABARAP-L2 remains dispersed in the cytosol (A). WT and S93/96A Δ C-terminal LC3C form autophagosomes that localize with HA-Parkin, whereas S93/96D Δ C-terminal LC3C remains dispersed in the cytosol (B). (C) SDS-PAGE and Western blot of HEK293T cell lysates transfected with HA-GABARAP-L2 Δ C-terminal WT or mutants. Cells were left untreated or treated with 40 µM CCCP for 3 hours to induce mitophagy.

**Supplementary Figure 6:**
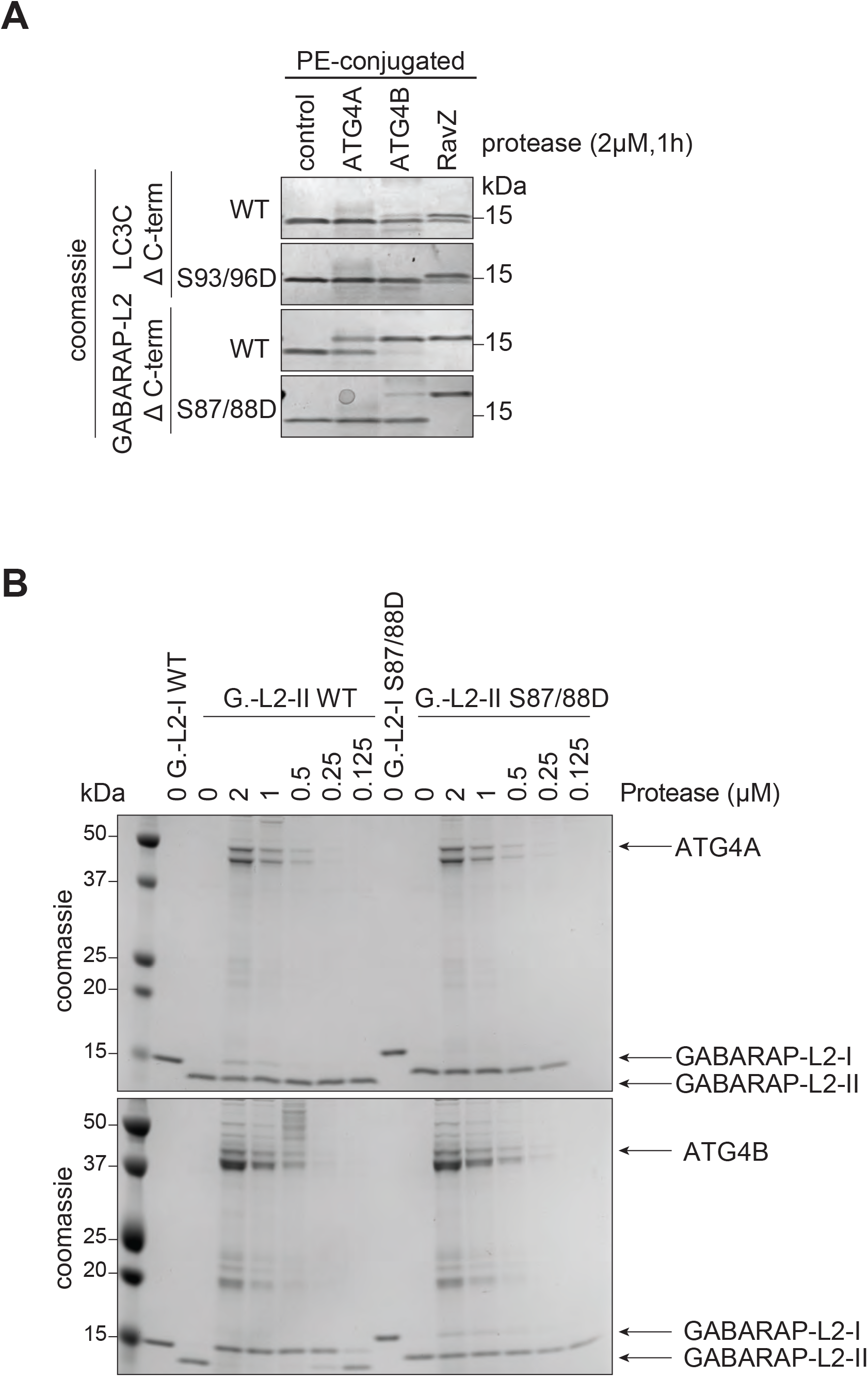
De-lipidation of GABARAP-L2 and LC3C by ATG4A, B and RavZ is dose dependent. (A) LC3C WT-, LC3C S93/96D-, GABARAP-L2 (G.-L2) WT- or GABARAP-L2 S87/88D-conjugated liposomes were treated or not with 2 µM ATG4A, ATG4B or RavZ for 1 hour at 37°C. Samples were then subjected to SDS-PAGE. (B) GABA-RAP-L2 WT- or GABARAP-L2 S87/88D-conjugated liposomes (G.-L2-II) were treated or not with different amounts (2-0.125 µM) of ATG4A or ATG4B for 1 hour at 37°C. Samples were then subjected to SDS-PAGE together with unconjugated GABARAP-L2 (G.-L2-I).

## Supporting Movie legends

### SM1: MD simulation of phosphorylated LC3C=ATG4B complex

First 200 ns of a 1158 ns trajectory showing the effect of phosphorylation of LC3C (red cartoon) on ATG4B interaction (grey surface). Phosphorylation at S93 and S96 (sticks) induces strong electrostatic effects that partially dislodge LC3C from the surface of ATG4B (first ~10-50 ns). The phosphorylated S93-PO_4_ and S96-PO_4_ detach from the ATG4B surface, opening a gap in the binding interface. The LC3C structure is distorted upon phosphorylation of S93 and S96. Its C-terminal tail retracts partially from the ATG4B active site. However, the ATG4B C-terminal LIR motif maintains stable interactions with the LIR binding pocket of LC3C (on the opposite face, Modeled).

### SM2: MD simulation of unphosphorylated LC3C=ATG4B complex

First 200 ns of a 1536 ns trajectory showing the dynamics of unphosphorylated LC3C (red cartoon) on ATG4B interaction (grey surface). S93 and S96 (sticks) are strongly bound to the ATG4B and buried in the interface. S93 and S96 of LC3C (red cartoon) mediate a series of polar contacts across the interface that provide specificity to ATG4B binding. S96 forms hydrogen-bonds with E350 of ATG4B. S93 is part of a network of polar contacts involving Y239, R229, N172 of ATG4B and Y86 of LC3C. Interactions across the extended interface allow stable sequestering of the C-terminal tail of LC3C close to the ATG4B active site throughout the simulation.

### SM3: MD simulation of full-length unphosphorylated LC3C_1-147_

First 500 ns from a representative trajectory (run 1 in Figure S3A) of full-length LC3C_1-147_ (red) showing large fluctuations of the unprocessed C-terminal tail. The large fluctuations of the C-terminal tail prevent R134 (sticks) to approach the phosphorylation sites S93 and S96 (sticks). The ATG4B binding site (grey surface) displays minimal fluctuations due to C-terminal tail dynamics.

### SM4: MD simulation of S93-PO_4_-LC3C_1-147_

First 500 ns from an example trajectory (run 3 in Figure S3B) of S93-PO_4_-LC3C_1-147_ (blue) showing the formation of a stable salt bridge between S93-PO_4_ and R134 (sticks). The movie shows the initial fluctuations of C-terminal tail enabling contact between S93-PO_4_ and R134 followed by rearrangement of the loop to form a stable salt bridge between. The ATG4B binding site (grey surface) is perturbed by the formation of salt bridge.

### SM5: MD simulation of S96-PO_4_-LC3C_1-147_

First 500 ns from an example trajectory (run 5 in Figure S3C) of S96-PO_4_-LC3C_1-147_ (green) showing the quick formation and disassociation of a salt bridge between S96-PO_4_ and R134 (sticks; within 80-100 ns). Subsequently, due to loop-rearrangement in the C-terminal tail, the R134 approach to S96-PO_4_ is restricted, leading to weak interactions in close-proximity (~0.5-0.6 nm). The ATG4B binding site (grey surface) is perturbed upon phosphorylation.

**Table S1:** Molecular dynamics simulations of LC3C and LC3C-ATG4B complexes. The table lists the different simulations of LC3C_1-147_ and of LC3C-ATG4B complexes, including the phosphorylation state, the number of runs, the total simulation time, the number of ions and water molecules, the total number of atoms, and the number of salt-bridge formation events between phosphorylated S93 or S96 and R134. In the column “interface interactions,” we give a qualitative assessment on the preservation of the interface structure and interactions in the LC3C-ATG4B complex simulations.

| LC3C | No of replicates | Total Simulation time (μs) | Ions (Na+/Cl-) | Tip3p Water | Total number of atoms | Salt-bridge formation events |
| --- | --- | --- | --- | --- | --- | --- |
| WT | 6 | 7.380 | 47/51 | 17078 | 53723 | 0 |
| S93-PO<sub>4</sub> | 6 | 7.155 | 48/50 | 17057 | 53663 | 4 |
| S96-PO<sub>4</sub> | 6 | 7.420 | 48/50 | 17101 | 53795 | 2 |
| S93/S93D | 6 | 7.181 | 60/62 | 21127 | 65896 | - |
| LC3C-ATG4B Complex | No of replicates | Simulation time (μs) | Ions (Na+/Cl-) | Tip3p Water | Total number of atoms | Interface interactions |
| WT | 1 | 1.475 | 106/91 | 30583 | 100457 | ++ |
| WT + LIR | 1 | 1.535 | 179/164 | 56164 | 177357 | +++ |
| S93/S96-PO<sub>4</sub><br>LC3C + LIR | 1 | 1.158 | 110/89 | 31711 | 103863 | — — |

**Table S2:** Binding free energy computations for LC3C-ATG4B complexes. The table lists different energetic contributions (mean ± s.d.) to the binding of LC3C and ATG4B in different phosphorylation states and with modeled LIR-WXXL interactions. The different binding energy contributions were computed using the MM-PBSA approach implemented in *g_mmpbsa* (see Methods) from MD simulations of LC3C-ATG4B complexes. The non-bonded energy terms (van der Waals and electrostatic) contribute significantly to the molecular-mechanics interaction energy of the complex, whereas changes in the bonded terms (bond-length, angle, and dihedral terms) do not contribute significantly to the interaction energy during complex formation.

| System/<br>Energy terms<br>[kJ/mol] | LC3C-ATG4B<br>(unphosphorylated) | LC3C-ATG4B<br>(unphosphorylated +<br>LIR) | LC3C-ATG4B<br>(S93/S96-PO<sub>4</sub> + LIR) |
| --- | --- | --- | --- |
| **van der Waals** | -1036.2 ± 110.2 | -985.9 ± 91.6 | -970.1 ± 128.4 |
| **Electrostatic** | -4652.7 ± 536.4 | -4495.8 ± 315.7 | -1610.2 ± 293.4 |
| **Polar solvation** | 2940.0 ± 406.0 | 2883.0 ± 339.2 | 2672.4 ± 311.0 |
| **SASA** | -136.2 ± 14.5 | -134.8 ± 10.2 | -125.2 ± 14.0 |
| **Total binding energy** | -2885.1 ± 305.2 | -2733.5 ± 204.0 | -33.1 ± 202.4 |
| **$\Delta\Delta G$** | - | 151.6 ± 367.1 | 2851.9 ± 366.2 |

